# Dissecting the Predictive Accuracy of Polygenic Indexes for Behavioral Phenotypes Across Genetic Ancestries

**DOI:** 10.1101/2025.09.11.675704

**Authors:** Robel Alemu, Alexander S. Young, Daniel J. Benjamin, Patrick Turley, Aysu Okbay

## Abstract

Polygenic indexes (PGIs) trained on samples of European genetic ancestries often lose substantial predictive power when applied to non-European ancestries. While this portability problem is well recognized, its manifestation in behavioral and social traits remains understudied, and the factors driving this accuracy loss warrant more comprehensive analysis. Using data from the UK Biobank and Health and Retirement Study, we conduct a systematic analysis of PGI portability for 52 health-related, behavioral, and social phenotypes. We advance prior literature by using genome-wide PGIs, assessing cross-ancestry heritability differences, and comparing the performance of PGIs based on standard versus family-based GWAS. Our findings confirm systematic reductions in PGI predictive power for non-European ancestries—with relative accuracy being lowest in African (24%), followed by East Asian (37%) and South Asian (51%) genetic ancestries. We also find that biologically proximal traits exhibit greater portability than behavioral and social traits. We show that the relative importance of factors underlying reduced portability varies across traits and ancestries: in African ancestries, linkage disequilibrium and allele frequency differences explain most of the loss (82%), compared with smaller contributions in East (34%) and South Asian (25%) ancestries. Finally, we find that family-based GWAS PGIs can modestly improve portability for select traits, such as BMI in African ancestry, suggesting that part of the portability gap may reflect population-specific confounds in standard PGIs.

## 1 Introduction

Genome-wide association studies (GWASs) have primarily been conducted on individuals of European (EUR) genetic ancestries, resulting in a lack of genetic diversity in GWAS datasets ^1–3^. Recent estimates indicate that less than 14 percent of all GWAS participants are from non-EUR genetic ancestries, a disparity with significant implications for the generalizability of genetic research ^4^. Although recent efforts have increased the representation of Asian ancestry groups in genetic studies, African, Latin American, and other populations remain largely underrepresented, contributing to persistent data imbalances ^5–7^. These imbalances impact the predictive accuracy of polygenic indexes (PGIs, also called polygenic scores). Specifically, the predictive accuracy of a PGI has been shown to be substantially smaller when applied to samples with a different genetic ancestry than the GWAS discovery sample ^2,8–10^. Martin *et al.* ^2^ demonstrated that PGIs trained on EUR-genetic-ancestry discovery samples show a predictive accuracy reduction of approximately 37%, 50%, and 78% for individuals of South Asian (SAS), East Asian (EAS), and African (AFR) genetic ancestries, respectively, across 17 biologically-proximal traits. Privé *et al.* ^11^ extended these findings by analyzing 245 traits, showing that PGI predictive accuracy not only diminishes across ancestries but also varies within continental ancestries as a function of genetic distance from the training population, a finding confirmed by Ding *et al.* ^12^ for over 80 phenotypes in the genetically diverse Los Angeles Biobank (ATLAS).

The limited portability of PGIs across genetic ancestries poses a significant challenge, potentially exacerbating existing disparities as the benefits of genetic research may not be equitably distributed among populations ^2,3,13–16^. As an initial step toward addressing this challenge, it is essential to understand the factors contributing to the reduced predictive accuracy of PGIs across different ancestries. Previous studies have suggested that factors such as cross-population differences in minor allele frequency (MAF), linkage disequilibrium (LD) between causal and tagging SNPs ^8^, SNP heritability, and variability in causal SNP effect sizes (due to gene-environment interactions or population-specific causal variants) ^17–20^ may contribute to the limited portability of PGIs. To quantify the contribution of LD and MAF differences to the cross-ancestry relative accuracy (RA) of PGIs, Wang *et al.* ^8^ developed a theoretical framework which they then applied to 8 biologically proximal phenotypes in UK Biobank. In this study, we extend Wang *et al.*’s findings by investigating the predictive accuracy of PGIs across ancestries for 54 health-related, behavioral, social, and cognitive phenotypes in two cohorts. Since social and behavioral phenotypes are likely to be more strongly influenced by complex gene-environment interplay ^21–24^, we anticipate that PGIs for these traits may exhibit even weaker portability than those for more biological proximal traits. To test this, we compare the cross-ancestry relative accuracy of PGIs across different phenotype categories, assessing whether the decline in predictive accuracy is more pronounced for social and behavioral traits. We further investigate whether the contribution of LD and MAF differences to loss of predictive accuracy differs across phenotype categories.

In addition to including a wide range of phenotypes, we extend Wang *et al.*’s analyses in several ways. First, we use PGIs based on weights for ∼ 2.9 million SNPs adjusted for LD using the SBayesR methodology ^25^ as opposed to unadjusted weights for only genome-wide significant SNPs. Second, our weights come from the Social Science Genetic Association Consortium’s Polygenic Index Repository ^26^ and are based on the largest available GWAS samples for most traits. This allows us to analyze traits for which the UK Biobank subsample used by Wang *et al.* would not be a sufficiently large discovery sample. Thirdly, we extend the predictive framework by investigating how variability in SNP heritability across ancestries influences relative accuracy, rather than assuming constant heritability as in previous studies. These refinements allow for a more comprehensive understanding of the factors affecting PGI portability for complex traits.

A fourth extension considers how potential confounds in PGIs based on between-family GWAS SNP weights, hereafter *standard PGIs*, affect their cross-ancestry portability. Between-family GWAS SNP weights can be biased due to passive gene-environment correlation, including population stratification and indirect genetic effects from relatives, and assortative mating ^27–30^. These biases can influence PGI predictive accuracy ^31–34^ and if gene-environment correlations and assortative mating patterns differ across ancestries, they can also affect relative accuracy ^33^. To assess this possibility, we also analyze PGIs based on family-based GWAS SNP weights, –*fGWAS PGIs*– which mitigate these biases by leveraging the random segregation of alleles within families. Comparing the cross-ancestry relative accuracy of standard and fGWAS PGIs allows us to test whether such confounds differ by ancestry and, in turn, to better understand the factors limiting PGI portability.

Finally, we extend our analyses from the UK Biobank (UKB) to the Health and Retirement Study (HRS). This cross-cohort comparison allows us to clarify whether conclusions regarding PGI portability across diverse ancestries hold under different demographic and environmental contexts. The two cohorts differ substantially in their design and composition. UKB recruited a large sample of middle-aged adults (40-69 years) who were, on average, healthier and more affluent than the general UK population, a common feature of volunteer-based cohorts ^35,36^. In contrast, HRS was designed as a nationally representative sample of US adults over the age of 50, capturing a wider range of health and socioeconomic circumstances that more closely reflect the general population approaching retirement ^37^.

In what follows, we first present the relative accuracy of PGIs in non-European genetic ancestries, then examine several factors that may underlie their reduced predictive power: (i) cross-ancestry differences in LD and MAF, (ii) differences in heritability, and (iii) variation in gene–environment correlation and assortative mating. We conclude with a discussion of the implications of our findings.

## 2 Results

### 2.1 Cross-Ancestry Predictive Accuracy of PGIs

We computed standard PGIs for 54 phenotypes across UKB and HRS (34 phenotypes are present in both cohorts, 16 are unique to UKB, and 4 are unique to HRS), while fGWAS PGIs were available for a subset of 24 of these UKB phenotypes (Supplementary Tables 1, 2 and 9). To assign individuals to one of four genetic ancestries—European (EUR), South Asian (SAS), East Asian (EAS), or African (AFR)—we first estimated principal component (PC) loadings from the 1000 Genomes Phase 3 reference panel ^38^. We then projected study participants onto this PC space and assigned ancestry by comparing each participant’s first 10 PCs to the mean values for each 1000 Genomes ancestry (Methods). In UKB, sample sizes were sufficiently large for all four ancestries (162,963 EUR, 11,413 SAS, 2,216 EAS, 9,494 AFR). In HRS, however, the SAS and EAS subsamples were too small to permit reliable analyses, so we limited our analyses to the EUR (12,774) and AFR (3,593) groups (Table 1). Following Wang *et al.* ^8^, we omit the AMR ancestry from our analyses because of their complex admixture patterns. For the standard PGIs, we use between-family GWAS weights for ∼ 2.9 million SNPs from the second release of Polygenic Index Repository ^26^. These weights are adjusted for LD using the SBayesR methodology ^25^. The fGWAS PGIs are based on family-based GWAS weights from Tan et al. ^39^ for HapMap3 ^40^ SNPs, adjusted for LD using PRS-CS ^41^. All weights are based on EUR-genetic-ancestry GWAS that excluded the target samples. Further details on PGI computation are provided in the Methods. (Section 4.7).

**Table 1:**
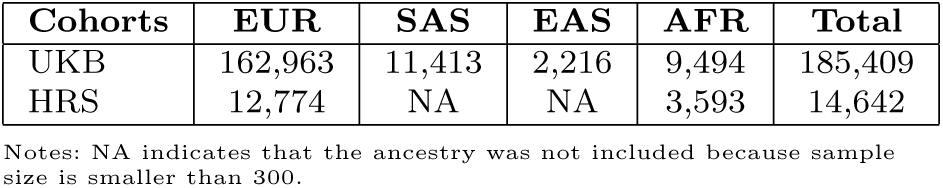
Number of genotyped samples by genetic ancestry and study cohorts.

| Cohorts | EUR | SAS | EAS | AFR | Total |
| --- | --- | --- | --- | --- | --- |
| UKB | 162,963 | 11,413 | 2,216 | 9,494 | 185,409 |
| HRS | 12,774 | NA | NA | 3,593 | 14,642 |

Notes: NA indicates that the ancestry was not included because sample size is smaller than 300.

We measure predictive accuracy using “incremental−*R*^2^”: the increase in the coefficient of determination (*R*^2^) when the PGI is added to a regression of the phenotype on the first 20 PCs of the genomic relatedness matrix (GRM) and also batch dummies in UKB. Prior to these regressions, we residualize the phenotypes on a third-degree polynomial in birth year, sex, and their interactions (Methods). To assess the loss in predictive accuracy when analyzing PGIs based on EUR-genetic-ancestry GWAS in non-EUR-genetic-ancestry samples, we compute the observed relative accuracy (*RA*_Obs_). Specifically, *i*- to-EUR relative accuracy (*RA*_obs_*^i^*) is defined as the ratio of the incremental *R*^2^ in *i*-ancestry to that in EUR-ancestry samples within the target cohorts (UKB and HRS):

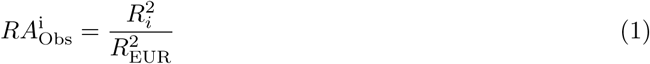

where, *R*_EUR_^2^ and *R_i_*^2^ denote the incremental *R*^2^ of the PGI for EUR and *i*-genetic-ancestry populations, respectively, with *i* ∈ {AFR, SAS, EAS}. To obtain the 95% confidence intervals (CIs) for incremental *R*^2^ and *RA*_Obs_, we bootstrap with 1000 replications and take the 2.5th and 97.5th percentiles as the lower and upper bounds, respectively.

Estimates of the predictive accuracy of standard PGIs across genetic ancestries in our target samples are presented in Supplementary Tables 1-2. Figures 1 and S1 and Supplementary Tables 4-5 show the relative accuracies. Across all 47 phenotypes in UKB, the average standard PGI relative predictive accuracy (*RA*_Obs_) is lowest in the African genetic ancestry group (28.03%, S.E. = 5.5), followed by East Asian (31.48%, S.E. = 3.4) and South Asian genetic ancestries (54.18%, S.E. = 4.7). In HRS, the average *RA*_Obs_^AFR^ is 23.31% (S.E. =6.8) across 33 phenotypes. Supplementary Figures S2 and S3 provide a detailed comparison of the incremental *R*^2^ estimates between EUR and non-EUR genetic ancestries for each phenotype in the UKB and HRS cohorts, respectively. Overall, our findings are consistent with previous studies by Martin *et al.* ^2^ and Wang *et al.* ^8^, which assessed the relative predictive accuracy of standard PGIs for a smaller set of biologically proximal traits (17 and 8 biologically proximal traits, respectively) in the UKB cohort. Expanding the analysis to 46 phenotypes, we find a smaller average relative predictive accuracy for standard PGIs in East Asian and South Asian ancestries compared to Martin *et al.*, while the results for African ancestry are comparable.

**Fig. 1:**
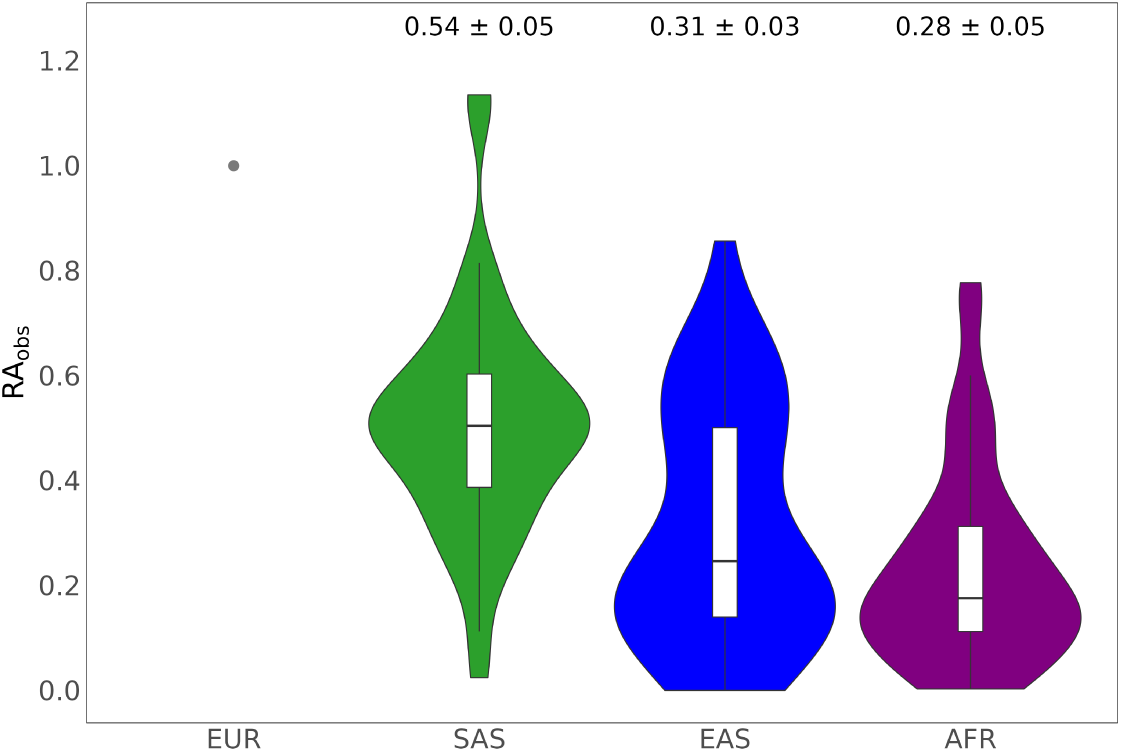
Violin plot of the relative predictive accuracy of standard PGIs for 46 phenotypes in the SAS, EAS, and AFR genetic ancestry samples from the UKB cohort. The violin plots illustrate the distribution and density of observed relative predictive accuracy for PGIs across different genetic ancestry groups, shown in comparison to European ancestry (marked by the gray dot). Box plots within the violin plots display the first, second (median), and third quartiles of the observed relative predictive accuracies (*RA*_Obs_). Labels above each plot provide the mean *RA*_Obs_ for each non-European ancestry group, calculated across 46 phenotypes, along with their standard errors, estimated using a leave-one-phenotype-out jackknife approach (see Supplementary Note 5.1.2). Incremental-*R*^2^ and *RA*_Obs_ values for each phenotype-ancestry pair are available in Supplementary Tables 1 and 3. For clarity of visualization, schizophrenia is excluded from this figure due to highly imprecise estimates (large standard errors) that would otherwise distort the scale (estimates are, however, reported in supplementary figures. Genetic ancestries: EUR = European, SAS = South Asian, EAS = East Asian, AFR = African.

To better understand differences in standard PGI relative accuracy across traits, we disaggregate the results by phenotype categories. In the UKB, the relative accuracy (*RA*_Obs_) is consistently highest in South Asian (SAS) ancestry, followed by East Asian (EAS) and African (AFR) ancestries (Figure 2). The highest mean *RA*_Obs_ for SAS (0.92; 95% CI: 0.12, 1.72) and AFR (0.88; 95% CI: −0.16, 1.92) ancestries is observed for psychiatric traits in UKB, driven mainly by the schizophrenia PGI being more predictive in these ancestries which have a higher case prevalence compared to EUR ancestry. In EAS, the highest mean *RA*_Obs_ is observed for blood biomarkers (0.62; 95% CI: 0.58, 0.66), which also rank second for SAS (0.65; 95% CI: 0.59, 0.71) and AFR (0.36; 95% CI: 0.26, 0.46). By contrast, the lowest *RA*_Obs_ in all ancestries is found for cognition and education—AFR (0.12; 95% CI: 0.10, 0.14), SAS (0.43; 95% CI: 0.33, 0.53), and EAS (0.15; 95% CI: 0.11, 0.19).

**Fig. 2:**
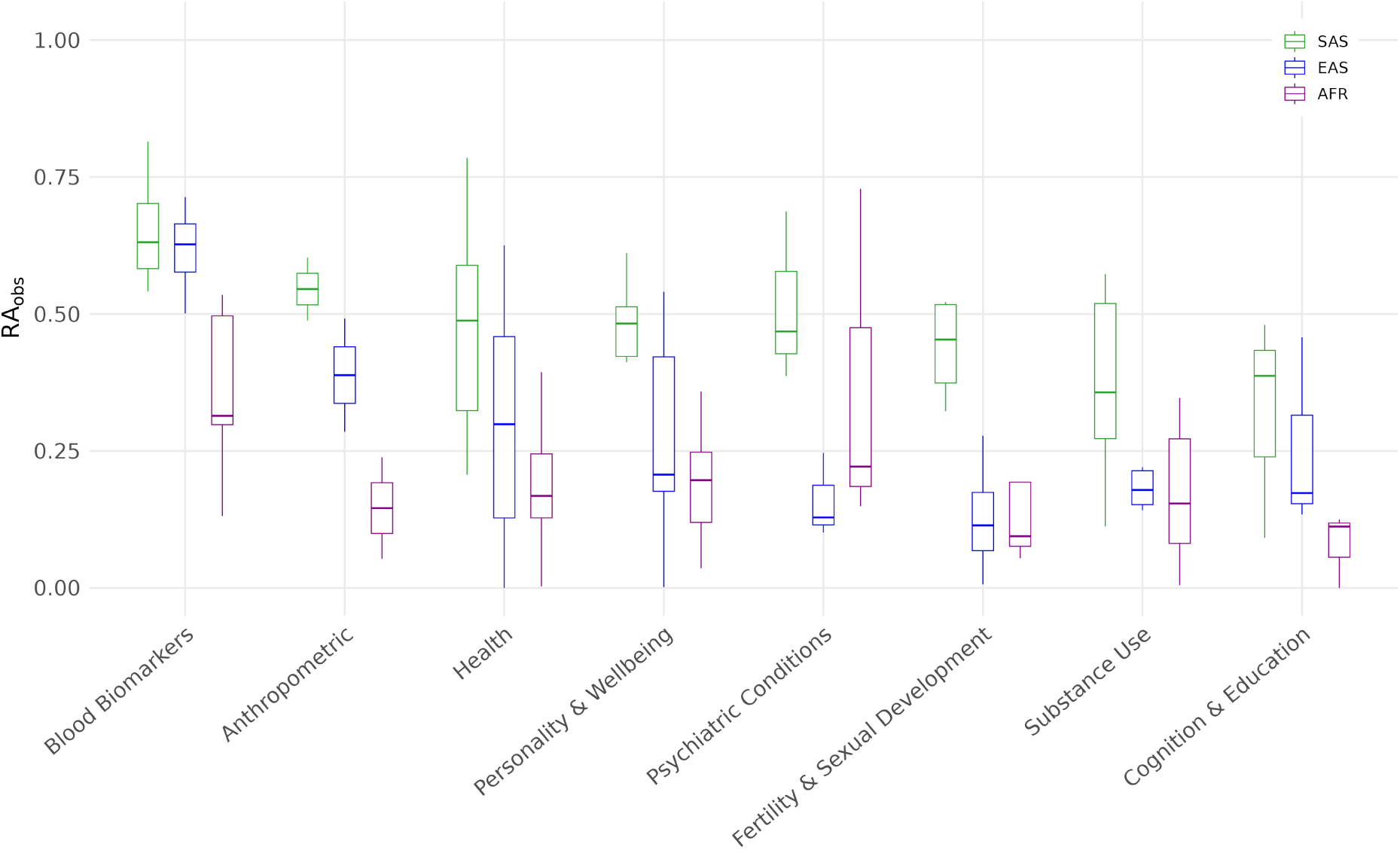
Observed relative accuracy of PGIs disaggregated by phenotype category. The box plots show the first, second (median), and third quartiles of *RA*_Obs_ averaged across phenotypes within each phenotype category by genetic ancestry. The list of individual phenotypes that constitute each category and their respective *RA*_Obs_ is presented in Supplementary Tables 4 (UKB) and 5 (HRS). *Note: Within the psychiatric conditions category, schizophrenia (SCZ) is excluded from the plot due to imprecise estimation that would distort the scale; it is, however, reported in Supplementary Table 4*.

In HRS, the highest mean *RA*_Obs_ is observed for substance use traits (0.57; 95% CI : −0.12, 1.26), largely driven by a greater predictive power of the PGI for alcohol misuse in AFR relative to EUR (Supplementary Table 5). This is followed by blood biomarkers (0.27; 95% CI : 0.19, 0.385, while the lowest predictive accuracy is observed for psychiatric conditions (0.08; 95% CI : −0.04, 0.20) and for fertility and sexual development (0.11; 95% CI : 0.00, 1.57)..

To formally summarize the extent of these differences in UKB, we applied Welch’s ANOVA to test whether mean *RA*_Obs_ differs across ancestry groups within each category. While we expect some level of cross-ancestry variation for all categories, we wanted to see if these differences are sufficiently large relative to within-category variability . We observed nominally significant differences in blood biomarkers, health, and fertility and sexual development. After Benjamini–Hochberg correction ^42^ (FDR = 5%), blood biomarkers and health remained significant (adjusted *p* = 0.007 and 0.045), whereas fertility and sexual development did not (Supplementary Table 6).

Our results are broadly consistent with prior studies, including Wang *et al.* ^8^ and Martin *et al.* ^2^, in that the average *RA*_Obs_ declines with increasing genetic distance from the reference group (EUR): highest in SAS, followed by EAS, and lowest in AFR ancestries. However, for some phenotypes, our estimates of *RA*_Obs_ differ from those reported by previous studies. For example, we find considerably higher *RA*_Obs_ for LDL-cholesterol in AFR and EAS but lower in SAS relative to Wang *et al.* ^8^. For height, we observe consistently higher *RA*_Obs_ across all non-EUR ancestries, while for BMI, our estimates are lower across the board. In the case of HDL-cholesterol, our estimate exceeds that of Wang *et al.* ^8^ only in AFR. For asthma, we observe uniformly lower *RA*_Obs_ across all ancestries. These divergences may be due to methodological differences, including the use of genome-wide SNPs (as opposed to genome-wide significant SNPs) and SNP weights derived from larger discovery samples in our study. For phenotypes whose GWS SNPs exhibit larger cross-ancestry MAF and LD differences compared to more weakly associated SNPs, all else being equal, we would expect our *RA*_Obs_ estimates to be larger. To provide a glimpse into the genetic architecture of the traits analyzed, Supplementary Figures S8–S17 display the effect sizes and ancestry-specific minor allele frequencies (MAF) of top associated SNPs, disaggregated by phenotype category.

Comparisons with Martin *et al.* ^2^ are more limited due to smaller phenotype overlap, but our estimates are lower for BMI and higher for both height and educational attainment in AFR genetic ancestry. Here, it is important to note that these cross-study differences should only be interpreted descriptively, as they are not based on formal statistical tests. Such tests would require joint re-estimation of both sets of *RA*_Obs_ values on the same individual-level data to properly account for sampling covariance. Collectively, however, these contrasts underscore how PGI construction choices and differences in discovery samples can influence cross-ancestry PGI portability, even when general patterns remain aligned across studies.

To better understand how potential confounds in standard PGIs influence cross-ancestry prediction in UKB, we compared their performance to that of fGWAS PGIs, which are constructed using weights that are not biased by indirect genetic effects from relatives, population stratification, and assortative mating. While these biases can inflate predictive accuracy in the discovery population, they may not translate well across divergent genetic backgrounds—potentially reducing cross-ancestry portability. We present the incremental *R*^2^ and *RA*_Obs_ for the fGWAS PGIs in Supplementary Tables 9 and 11, respectively.

Because the fGWAS have much smaller effective sample sizes, the incremental-*R*^2^ values from fGWAS PGIs are generally smaller. In many cases, the non-EUR incremental-*R*^2^ is statistically indistinguishable from zero (19/24 phenotypes in AFR, 8/24 in SAS, and 17/24 in EAS). Despite this, several traits exhibit comparable or higher *RA*_Obs_ in fGWAS PGIs relative to their standard GWAS counterparts. Most notably, the relative accuracy of the fGWAS PGI for BMI in AFR is significantly higher than that of the standard PGI, with *RA*_Obs_ = 0.34 (95% CI: 0.23, 0.46) for fGWAS versus 0.05 (95% CI: 0.03, 0.09) for the standard PGI; this difference remains statistically significant after multiple-testing correction (*P* = 0.001). We also observe nominally significant improvements in *RA*_Obs_ for BMI in SAS (*P* = 0.048), systolic blood pressure in AFR (*P* = 0.01), and ever smoking in EAS (*P* = 0.02), though these do not survive multiple testing correction (Figure 3). For the remaining traits, *RA*_Obs_ differences are not statistically distinguishable from zero, but the point estimates from fGWAS PGIs tend to be higher across most traits and ancestries (Supplementary Table 11). Details of the paired nonparametric bootstrap test for differences in *RA*_Obs_ between the two approaches (and the multiple-testing correction) are provided in Supplementary Note, Section 5.2.1.

**Fig. 3:**
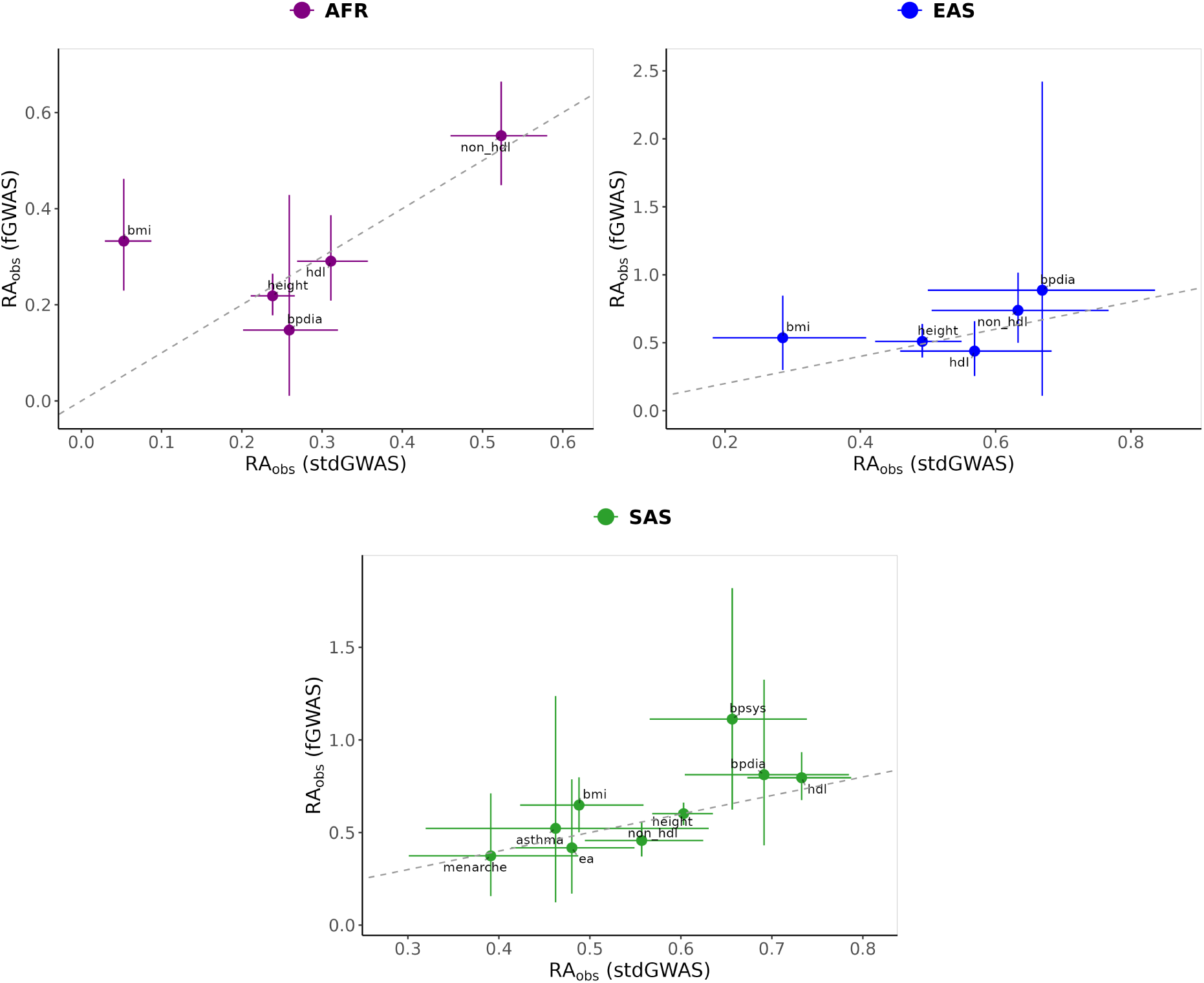
Comparison of observed relative accuracy (*RA*_Obs_) for fGWAS and standard GWAS PGIs across non-EUR Ancestries in UKB. Panels show *RA*_Obs_ values estimated using fGWAS-based PGIs on the y-axis versus those estimated using standard GWAS PGIs on the x-axis, with separate panels for each ancestry group: AFR (left), EAS (middle), and SAS (bottom). Each point represents a phenotype, with error bars indicating the 95% confidence intervals (CIs) for *RA*_Obs_ estimates. The diagonal dashed line marks the line of equivalence, where *RA*_Obs_ values for fGWAS and standard GWAS would be equal. Phenotypes with imprecisely estimated *RA*_Obs_ values were excluded from the figure for better visibility but remain included in Tables S4 and S10. Filtering was applied to remove phenotypes where (1) *RA*_Obs_ estimates were not significantly different from zero and (2) confidence interval widths exceeded three times the absolute point estimate.

### 2.2 Analysis of Factors Influencing PGI Predictive Accuracy

#### 2.2.1 Cross-ancestry LD and MAF differences

To evaluate the contribution of different factors to the loss of cross-ancestry PGI predictive accuracy, we have generalized the approach proposed by Wang *et al.* ^8^. Similar to Wang *et al.*, we use a model that expresses the predicted PGI relative accuracy as a function of cross-population differences in LD, MAF, SNP-based heritability, as well as the cross-population correlation of causal SNP effect sizes. Under this model, Wang *et al.*, derived the expected relative accuracy of PGIs (*RA*_Expected_) as:

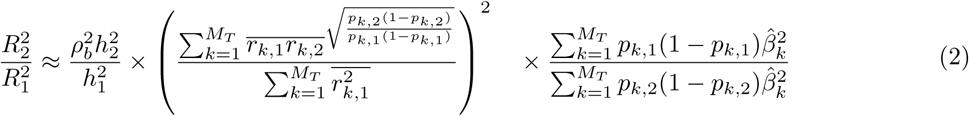

where *ρ_b_*^2^ is the correlation of causal SNP effect sizes between populations 1 and 2, *h_i_*^2^ is the SNP-based heritability in population *i* = {1, 2}, *p_k,i_* is the minor allele frequency (MAF) of PGI-SNP *k* in population 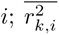 is the mean squared correlation of allele counts between PGI-SNP *k* and all “candidate causal SNPs” in LD with it in population 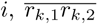 is the mean product of the correlation of allele counts between PGI-SNP *k* and all “candidate causal SNPs” in LD with it in populations 1 and 2, *β̂_k_*^2^ is the effect size of PGI-SNP *k* as estimated in the discovery GWAS, and *M_T_* is the number of SNPs in the PGI.

We deviate from Wang *et al.* in the set of PGI-SNPs and “candidate causal SNPs” used in Equation 2. Wang *et al.* focus on PGIs constructed using only independent genome-wide significant (GWS) SNPs, based on two key considerations: first, genome-wide significant SNPs are more likely to pinpoint causal variants compared to weaker associations; second, prior studies have shown that including sub-significant SNPs reduces the cross-ancestry relative accuracy of PGIs ^2,9,43,44^. Relying on results from a prior simulation study ^45^, they define “candidate causal SNPs” as SNPs in LD (*R*^2^ *>* 0.45) with GWS-SNPs within a 100 kb window. In contrast, we construct PGIs using Bayesian methodologies that incorporate a much larger set of common SNPs. For the main analyses, we use weights from the second release of PGI Repository, obtained using the SBayesR methodology with ∼ 2.9 million SNPs ^25^. For the fGWAS PGIs, we use the weights for ∼ 1.2 million HapMap3 SNPs ^40^ generated by Tan *et al.* ^39^ using PRS-CS ^41^. The reasons behind this are two-fold, concerning feasibility and practical implications. First, we consider a wider set of behavioral and health-related phenotypes in this study. These phenotypes are highly polygenic, with each SNP having a very small effect size, and current largest GWAS available for many of these phenotypes are only able to identify a handful of GWS SNPs. Consequently, relying solely on GWS-SNPs to calculate the predicted relative accuracy may yield imprecise results regarding the relevant cross-ancestry differences that drive the loss. Second, most PGI studies construct PGIs with a focus on maximizing predictive power for the phenotype. Although, as Wang et al. state, the accuracy of GWS-based PGIs will get closer to that of genome-wide PGI methodologies as GWAS sample sizes become larger, we are not there yet. Therefore, we wanted to generalize and assess Wang *et al.*’s model under practically more relevant conditions.

Including more than a million SNPs in the PGIs irrespective of their association p-values leaves us with the challenge of defining “candidate causal SNPs”. We need a set of SNPs that would approximate the genome-wide level expected predictive accuracy of PGIs. Because the PGI-SNPs are not selected based on their p-values and are not pairwise independent, we cannot assume that causal SNPs are located within a 100kb window and are in LD with the PGI-SNPs. Therefore, we decided to use Wang *et al.*’s approach of defining candidate causal SNPs in relation to the most significant independent SNPs, but relaxing the p-value cutoff to consider more than only GWS SNPs. The challenge then becomes including enough SNPs to correspond to the predictive power of genome-wide PGIs while keeping the computational burden at a manageable level. In order to gauge how the expected relative accuracy changes in relation to the number of candidate causal SNPs included in the model, we generated three different candidate causal SNP sets for each phenotype based on the top 100, 1,000, and 10,000 independent SNPs by p-value (Section 4.8). We then expanded each set to include any SNPs in LD (*R*^2^ *>* 0.45) within 100 kb of these top SNPs. Using Equation 2, we computed the expected relative accuracy of the PGIs based on each candidate causal SNP set, fixing 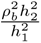 at unity. For the fGWAS PGIs, we repeated the procedure selecting top SNPs based on fGWAS p-values (Methods).

Our evaluations reveal that the expected relative accuracy (*RA*_Expected_), modeled based on cross-ancestry differences in MAF and LD, is generally stable after the top 1,000 SNPs for most phenotypes, whether computed with standard GWAS or fGWAS candidate SNP sets (Figure 4, Supplementary Figures S4-S6). Using the top 10,000 SNPs provides the most precise estimates for both approaches; thus, we use this set as a proxy for genome-wide expectations in subsequent analyses. For the standard GWAS-based candidate causal SNPs, the average *RA*_Expected_ across 54 phenotypes was 36% for AFR, 79% for EAS, and 87% for SAS genetic ancestry. The fGWAS-based SNPs yielded similar averages for SAS (87%) and EAS (77%), with a modest increase for AFR (39%). Ancestry-specific *RA*_Expected_ values estimated based on the top 10,000 candidate causal standard GWAS-based SNPs for individual phenotypes are provided in Supplementary Table 3, while those based on fGWAS SNPs are shown in Supplementary Table 10.

**Fig. 4:**
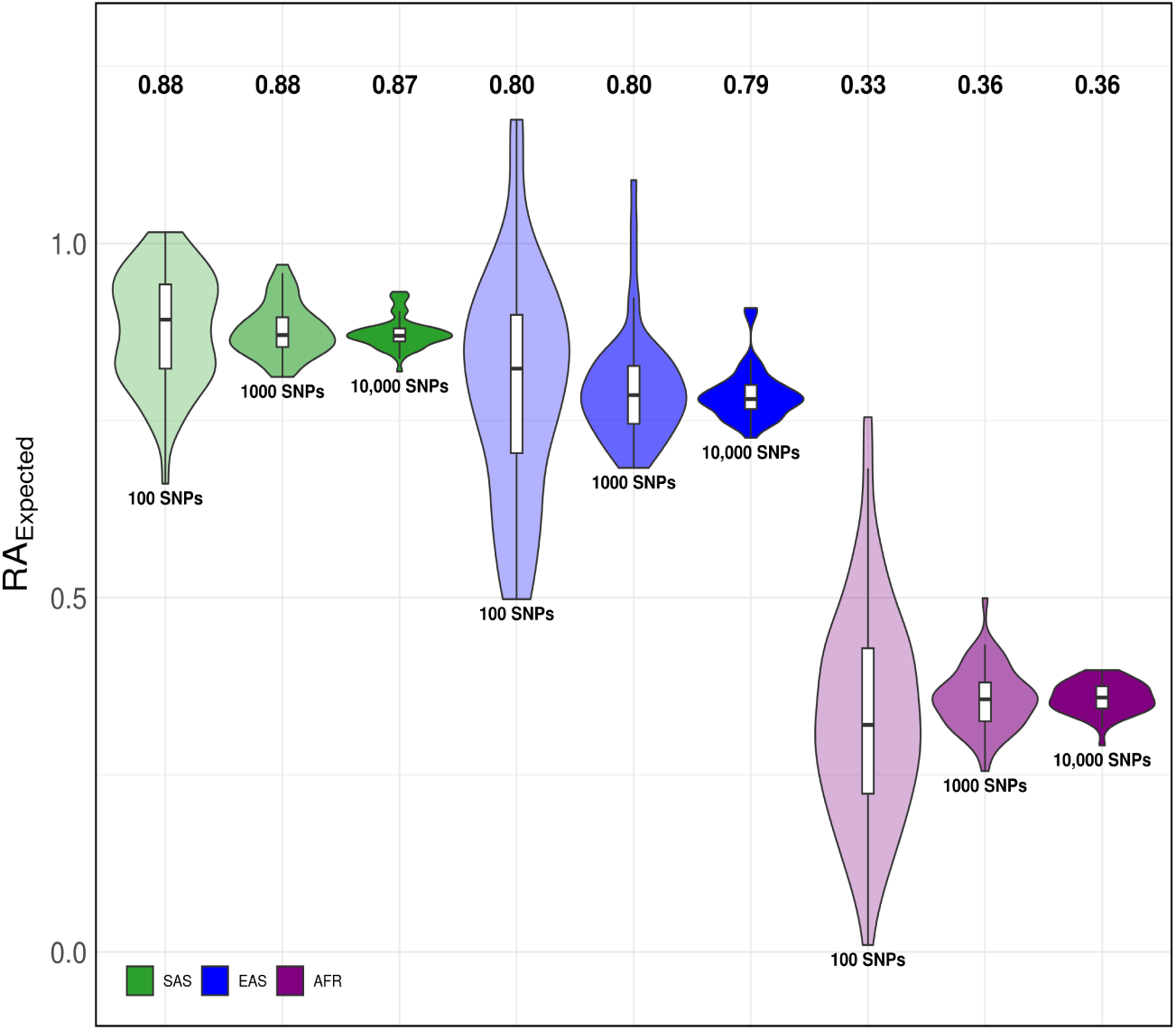
Average expected relative predictive accuracy of PGIs by different candidate causal SNP sets in non-European ancestries based on standard GWAS summary statistic. Violin plots show the distribution of *RA*_Expected_ values across 54 phenotypes, with inner box plots indicating the first, second (median), and third quartiles of these values. Labels above each plot indicate the mean *RA*_Expected_ for each ancestry, averaged over the phenotypes.

After estimating *RA*_obs_ and *RA*_Expected_, we calculate the percentage loss in PGI predictive accuracy attributable to cross-ancestry differences in LD and MAF, denoted as *LoA*_(LD+MAF)_. We compute this measure only for phenotypes that exhibit a *RA*_obs_ statistically smaller than one, indicating meaningful shrinkage in the PGI’s predictive accuracy. For these phenotypes, *LoA*_(LD+MAF)_ is derived as the ratio of the expected loss to the observed loss in PGI accuracy, as formalized in Equation (3). A value of 100% indicates that differences in LD and MAF fully account for the reduction in predictive accuracy. Values below 100%—where *RA*_obs_ *< RA*_Expected_—suggest that the decline in predictive performance is larger than what LD and MAF differences alone would predict, pointing to additional contributing factors such as imperfect genetic correlations between the target and reference group (*ρ_b_*^2^ *<* 1) or lower heritability in the target population (*h*_2_^2^ *< h*_1_^2^). Conversely, values above 100%—where *RA*_obs_ *> RA*_Expected_—indicate that the observed loss is smaller than expected based on LD and MAF, implying that other factors may be partially compensating for the predicted reduction. One plausible explanation is that the heritability of the trait is higher in the target population than in the discovery cohort (*h*^2^ > *h*^2^), which would enhance the observed predictive accuracy beyond what is predicted from LD and MAF differences alone.

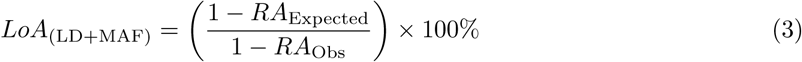

Our analysis of *standard GWAS* -based PGIs in the UKB cohort reveals substantial differences in the mean loss of predictive accuracy due to LD and MAF across non-EUR genetic ancestries and phenotype categories (Figure 5, Supplementary Table 4). The mean *LoA*_(LD+MAF)_ is highest in AFR, averaging 83.04% (S.E. = 2.9%) across 41 phenotypes, followed by EAS at 32.50% (S.E. = 2.3%) over 32 phenotypes, and SAS at 24.90% (S.E. = 1.9%) across 36 phenotypes. Breaking down by phenotype category, *LoA*_(LD+MAF)_ is most pronounced for blood biomarkers across all non-EUR ancestries: 106.05% (S.E. = 8.8%) in AFR, 47.08% (S.E. = 6.3%) in EAS, and 35.59% (S.E. = 6.8%) in SAS. In contrast, the lowest mean *LoA*_(LD+MAF)_ values are observed for fertility and sexual development traits in AFR (69.70%; S.E. = 1.1%) and EAS (26.25%; S.E. = 0.5%), and for substance use traits in SAS (17.94%; S.E. = 1.5%). Beyond these patterns, differences across the remaining categories appear modest, with anthropometric traits showing a slightly elevated mean *LoA*_(LD+MAF)_ in EAS and SAS (Figure S7).

**Fig. 5:**
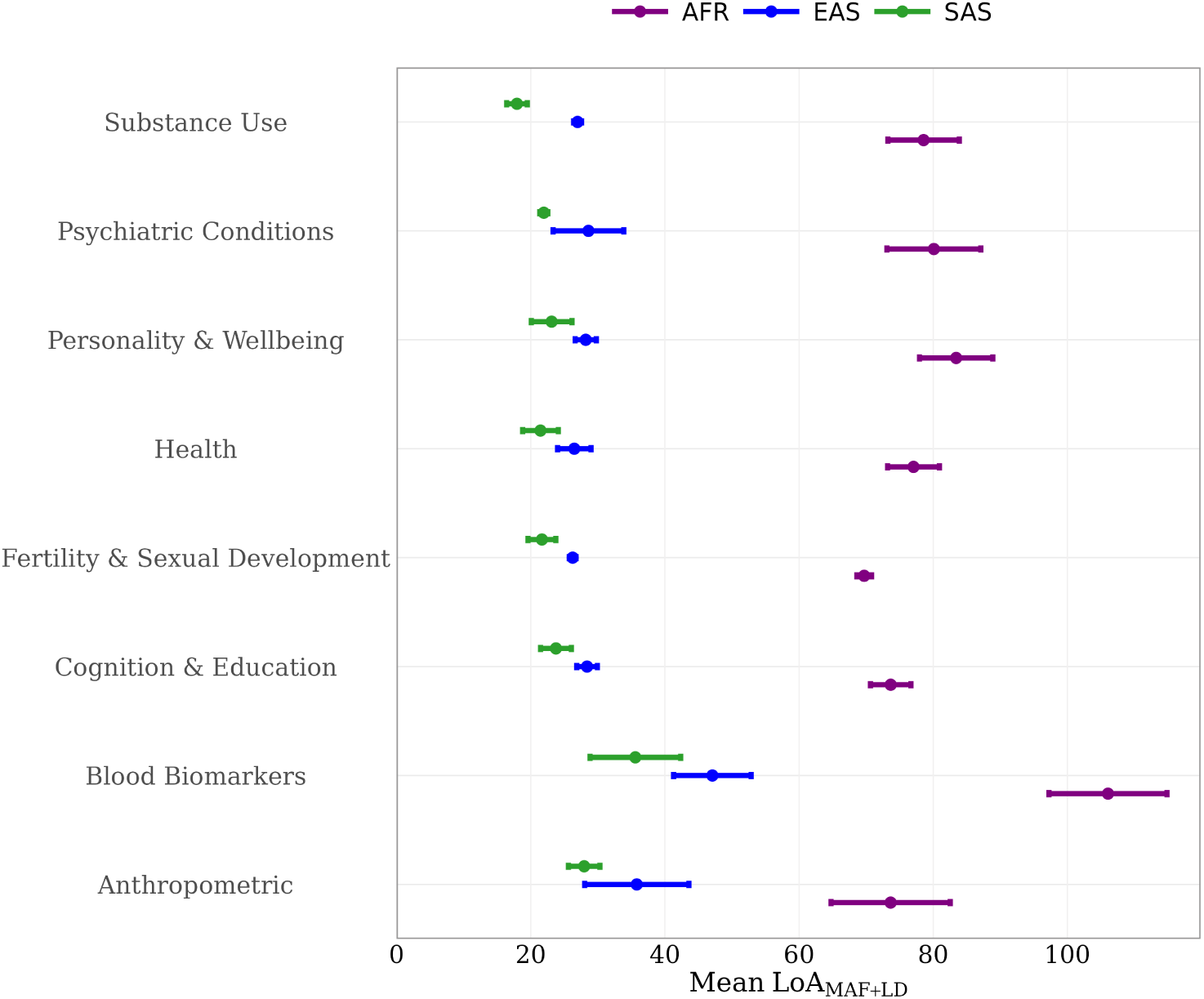
Contribution of MAF and LD to the Loss of PGI Predictive Accuracy Across Phenotype Categories in the UKB Cohort. This figure illustrates the average loss of predictive accuracy in *standard GWAS* -based PGIs attributable to MAF and LD in African (AFR), East Asian (EAS), and South Asian (SAS) genetic ancestry samples from the UKB across eight phenotype categories. The mean *LoA*_(LD+MAF)_ values are shown as point estimates, with error bars representing 95% confidence intervals calculated using a leave-one-phenotype-out jackknife approach. *LoA*_(LD+MAF)_ estimates for individual phenotypes within each category are provided in Supplementary Figure S7 and in Supplementary Table 4.

Zooming in on where *LoA*_(LD+MAF)_ is largest - blood biomarkers - we see pronounced heterogeneity within the category (Figure S7). In EAS, blood pressure phenotypes (systolic: 70.89%, S.E. = 6.1%; diastolic: 61.3%, S.E. = 7.3%; pulse: 56.41%, S.E. = 7.9), and HDL cholesterol (57.46%, S.E. = 7.7%) show outstandingly high *LoA*_(LD+MAF)_ values. In SAS, the largest *LoA*_(LD+MAF)_ is observed for triglycerides (71.69%, S.E. = 12.7%), followed by HDL cholesterol (47.91% (S.E. = 7.3%). In contrast, LDL and non-HDL cholesterol exhibit some of the lowest *LoA*_(LD+MAF)_ values in both EAS (LDL: 20.81%, S.E.=15.3; non-HDL: 24.80%, S.E.=17.3) and SAS (LDL: 15.80%, S.E. = 8.5; non-HDL: 15.42%, S.E. = 7.3) ancestries, which is to be expected given that LDL and non-HDL cholesterol are known to be more strongly influenced by lifestyle factors compared to HDL cholesterol ^46^. Interestingly, this pattern does not hold in AFR ancestry, where LDL (133.61%, S.E. = 6.05%) and non-HDL (126.41%, S.E. = 5.75%) cholesterol exhibit the highest *LoA*_(LD+MAF)_ values, with HDL cholesterol following a few phenotypes behind (102.70%, S.E.=3.4).

The lowest *LoA*_(LD+MAF)_ is observed for COPD, the leading cause of which is cigarette smoking, in both EAS (9.67%, S.E.=6.8) and SAS (9.92%, S.E.=5.0) ancestries. COPD is also one of the lowest for AFR (68.66%, S.E.=4.8). Prostate cancer has the lowest *LoA*_(LD+MAF)_ in AFR genetic ancestry (62.96%, S.E.=2.5), ranks second lowest in SAS (12.06%, S.E.=2.4) and fourth lowest in EAS (62.96%, S.E.=2.5) (Supplementary Table 4 and Figure S7).

For a small subset of traits, we observe *LoA*_(LD+MAF)_ values exceeding 100% only in AFR. The phenotypes with *LoA*_(LD+MAF)_ significantly above 100% (95% CI entirely *>*100%) are: Life Satisfaction—Family (111.75%; 95% CI: 105.15, 118.35), total cholesterol (126.41%; 95% CI: 115.14, 137.69), non-HDL cholesterol (133.61%; 95% CI: 121.75, 145.47), and LDL cholesterol (142.42%; 95% CI: 130.50, 154.34). Similarly, a few other phenotypes had point estimates above 100% but were not statistically greater than 100%—namely, smoking cessation (101.06%; 95% CI: 96.46, 105.65), HDL cholesterol (102.70%; 95% CI: 95.96, 109.43), and type-II diabetes (105.43%; 95% CI: 99.67, 111.18). Consistent with this pattern, Wang *et al.* also report *>*100% values for certain traits in AFR (e.g., LDL cholesterol: 124.90% with S.E. = 10.5%; asthma: 107.3% with S.E. = 27.0%). *LoA*_(LD+MAF)_ higher than *>*100%) indicates that factors beyond LD and MAF may be positively influencing the observed predictive accuracy of PGIs for these traits. In subsequent sections, we explore whether accounting for additional factors such as heritability differences or using fGWAS-based approaches to compute PGIs alters this scenario.

Overall, our findings are in line with Wang *et al.*. Although there are substantial differences in point estimates for some phenotypes such as asthma in AFR and BMI in EAS where our estimates are lower and type-II diabetes in AFR and height in EAS where our estimates are higher, these estimates are contained within the substantially wider 95% confidence intervals reported by Wang *et al.*. Exceptions to this are LDL cholesterol in EAS and SAS, and height in AFR and SAS ancestries. Our *LoA*_(LD+MAF)_ values for LDL cholesterol in EAS and SAS reported above are much lower compared to Wang *et al.*’s, who found 97.6% (S.E. = 23.8%) for EAS and 42.1% (S.E. = 2.7%) for SAS ancestries. For height, we find larger *LoA*_(LD+MAF)_ values for AFR (82.54%, S.E. = 1.8% vs. 71.50%, S.E. = 1.8%) and SAS (30.32%, S.E. = 2.4% vs. 23.6%, S.E. = 1.8%).

As with the differences in relative accuracy estimates, the differences in *LoA*_(LD+MAF)_ estimates between our study and Wang *et al.* likely stem from methodological distinctions, particularly in SNP selection and PGI methodology. Lupi *et al.* ^47^ demonstrate that RA_obs_ and LoA_LD+MAF_ vary substantially across the genome, even within a given ancestry. They show that certain genomic regions—referred to as “high portability” segments—exhibit consistently strong cross-ancestry predictive accuracy, including in populations such as AFR where genome-wide RA_obs_ is typically low. Lupi *et al.* also report that the LoA_LD+MAF_ estimates are markedly lower in these high portability regions across traits and ancestries. However, it is important to note that for certain phenotypes (height, LDL-and HDL-cholesterol), we observe substantially higher *RA*_Obs_ in our study relative to Wang *et al.*, while the *RA*_Expected_ based on MAF and LD differences is similar across studies. In fact, we find that for most phenotypes, *RA*_Expected_ due to MAF and LD differences alone is relatively stable after the top 1,000 SNPs, and inclusion of more high portability areas in the model would increase both *RA*_Obs_ and *RA*_Expected_. Several other explanations are possible. The simplest is that adjusting the SNP weights for LD improves the cross-ancestry portability of PGIs by getting closer to the causal effect sizes. Other explanations could include causal effects being more heterogeneous across ancestries for the top SNPs, or top SNPs not being representative of the whole genome in terms of the contribution of gene-environment correlation., e.g.

We next estimated standard PGI *LoA*_(LD+MAF)_ values for AFR ancestry in HRS. The highest *LoA*_(LD+MAF)_ in HRS is observed for COPD (104.39%, S.E. = 7.4%), followed by smoking cessation (97.17%, S.E. = 2.4%), systolic blood pressure (88.99%, S.E. = 1.6%), and height (88.48%, S.E. = 1.9%). The lowest *LoA*_(LD+MAF)_ estimates are for coronary artery disease (65.62%, S.E. = 1.7%) followed by a suit of behavioral phenotypes: subjective well-being (67.70%, S.E. = 1.4%), family satisfaction (71.26%, S.E. = 2.0%) and depressive symptoms (71.92%, S.E. = 1.4%). For a direct comparison with UKB, we plotted the *LoA*_(LD+MAF)_ estimates for AFR in both cohorts, displaying point estimates alongside their 95% confidence intervals (Figure 6). In UKB, *LoA*_(LD+MAF)_ estimates for migraine, subjective well-being, and depression are significantly higher than those in HRS after correcting for multiple testing using the Benjamini–Hochberg procedure ^42^ (FDR = 5%). Conversely, HRS shows significantly higher *LoA*_(LD+MAF)_ for phenotypes including BMI, educational attainment, drinks per week, height, cigarettes per day, and neuroticism. These differences suggest that the contribution of MAF and LD to PGI predictive accuracy may vary across cohorts, potentially reflecting differences in environmental exposures or sample characteristics. In some cases, substantial differences in phenotype definitions may also contribute to the observed discrepancies.

**Fig. 6:**
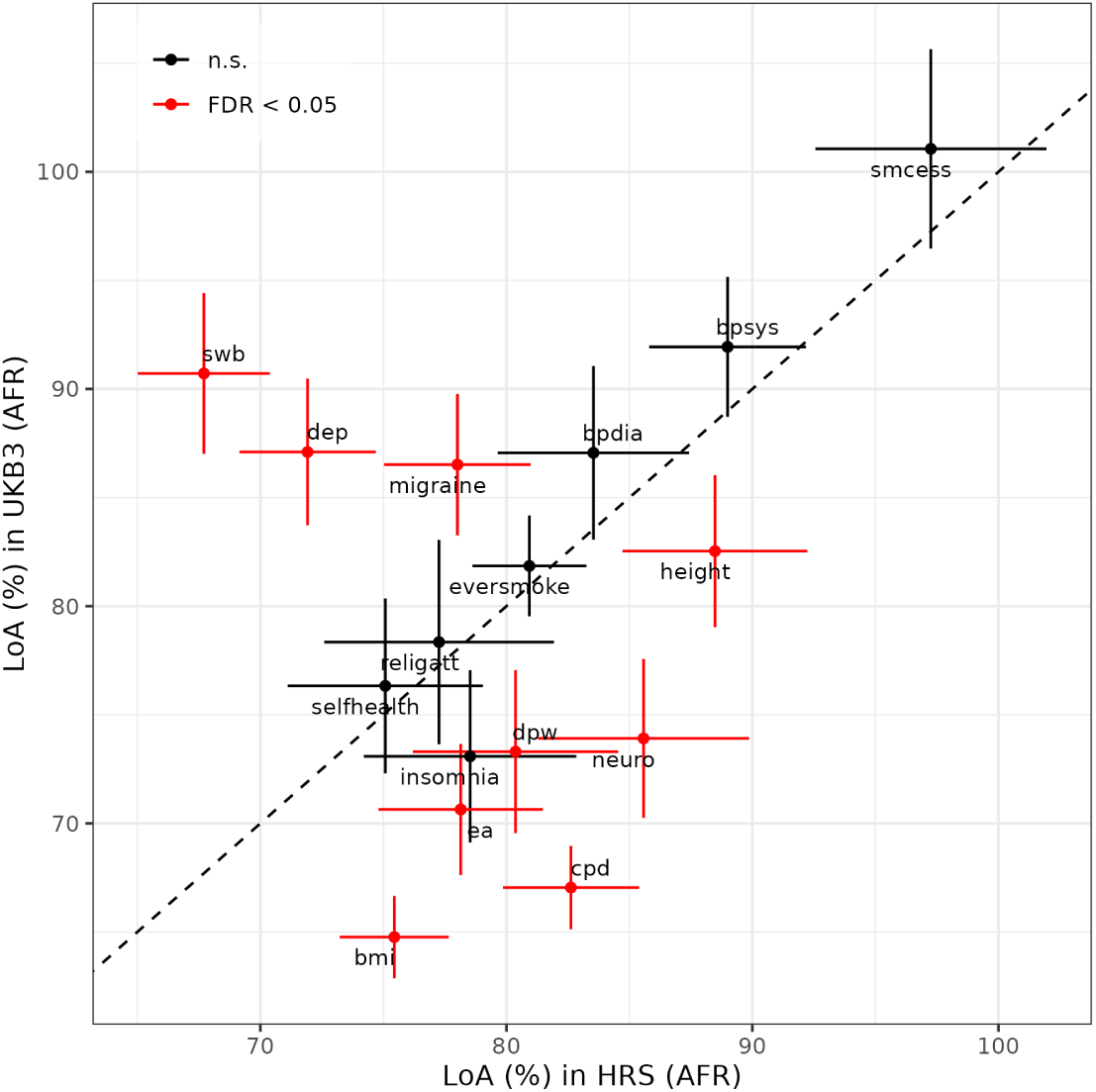
Comparison of the Contribution of MAF and LD to the Loss of PGI Predictive Accuracy Between HRS and UKB Cohorts in AFR Genetic Ancestry. Each point represents *LoA*_(LD+MAF)_ for a phenotype in HRS on the x-axis and UKB on the y-axis; error bars show 95% confidence intervals. The standard error of *LoA*_(LD+MAF)_ was obtained with the delta method (Supplementary Note 5.1.3) ^48^. The phenotype chronic obstructive pulmonary disease (COPD) was excluded from the plot as the *LoA*_(LD+MAF)_ was too imprecisely estimated (see Table S5). Statistical comparisons between cohorts were performed using a log-scale *Z*-test with FDR correction (for full details, see Supplementary Note 5.2.2). The dashed 45° line marks equality; red points indicate phenotypes whose *LoA*_(LD+MAF)_ differ significantly between cohorts at FDR *<* 0.05. n.s., not significant; FDR, false discovery rate.

#### 2.2.2 Cross-ancestry heritability differences

Next, we account for cross-ancestry differences in SNP-based heritability to further elucidate the combined effect of MAF, LD, and heritability (*h*^2^) in explaining the shrinkage of standard PGI predictive accuracy. To obtain the *LoA*_(LD+MAF+h_2 _)_, we adjust Equation (3) by multiplying the *RA*_Expected_ by the ratio of the SNP-based heritability estimates in the target non-EUR and EUR genetic-ancestry samples which we estimate using BOLT-REML (Methods). Given the relatively small non-EUR sample sizes in both UKB and HRS cohorts, heritability estimates for most traits were imprecise, yielding large standard errors (Supplementary Tables 1 and 2). This imprecision resulted in *LoA*_(LD+MAF+h_2 _)_ estimates that were statistically indistinguishable from zero for many traits, particularly for SAS and EAS genetic ancestries in UKB.

Within the AFR ancestry of UKB, the *LoA*_(LD+MAF+h_2 _)_ estimates for BMI, height, and HDL cholesterol were calculated with reasonable precision and were all higher than the corresponding *LoA*_(LD+MAF)_ values. For height, adding heritability to the LD+MAF model raised the share of variance explained in the loss of accuracy to nearly 100%, indicating that LD, MAF, and *h*^2^ together account for almost the entire reduction in PGI predictive power relative to EUR (Figure 7). In contrast, HDL and LDL cholesterol remained above 100% after accounting for heritability. This pattern suggests that factors beyond LD, MAF, and heritability contributing to the predictive accuracy of standard-GWAS PGIs such as passive gene–environment correlations or assortative mating may have inflated *RA*_obs_ in AFR ancestry, producing *RA*_obs_ *> RA*_Expected_ and, consequently, *LoA*_(LD+MAF+h_2 _)_ *>* 100%.

**Fig. 7:**
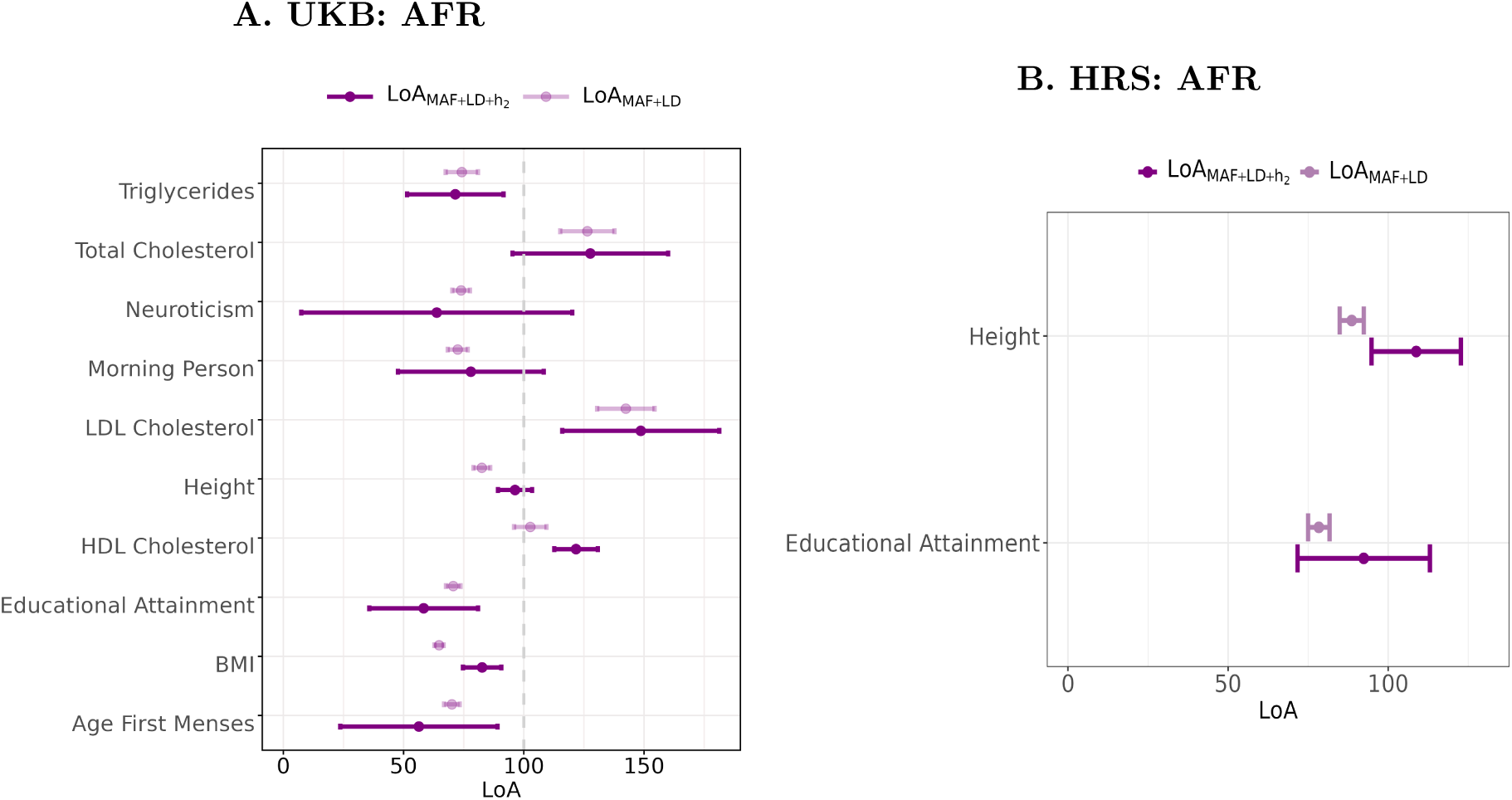
Comparison of the *LoA*_(LD+MAF)_ and *LoA*_(LD+MAF+h^2^)_ in AFR genetic-ancestry samples of the UKB and HRS cohorts. This figure displays two panels (A for UKB, B for HRS) comparing the *LoA*_(LD+MAF)_ with the *LoA*_(LD+MAF+h_2 _)_ for African genetic-ancestry samples. Each panel depicts the loss of predictive accuracy alongside 95% confidence intervals for selected phenotypes for which the estimates were statistically different from zero. Standard errors were calculated using the delta method ^48^ (Supplementary Note 5.1.3). Estimates for South Asian (SAS) and East Asian (EAS) ancestries are absent due to exceedingly large standard errors resulting from imprecisely estimated SNP-based heritabilities, making their *LoA*_(LD+MAF+h_2 _)_ estimates unreliable.

In the HRS cohort, we could estimate *LoA*_(LD+MAF+h_2 _)_ with acceptable precision only for height and educational attainment; for height, the pattern mirrored UKB, while educational attainment exhibited a similar but non-significant rise.

Overall, for phenotypes for which we were able to estimate AFR-genetic-ancestry SNP-based heritability relatively precisely, our findings suggest that accounting for cross-ancestry differences in SNP-based heritability largely addresses the remaining unexplained shrinkage in PGI predictive accuracy (Figure 7). However, the persistence of *LoA*_(LD+MAF+h_2 _)_ estimates exceeding 100% for traits such as HDL and LDL cholesterol motivates our subsequent analysis, where we assess whether PGIs derived from family-based GWAS which are not affected by passive gene–environment correlations and assortative mating can help resolve this discrepancy.

#### 2.2.3 *LoA*_(LD+MAF)_ for fGWAS based PGIs

Finally, we employ fGWAS-based PGIs to re-compute *LoA*_(LD+MAF)_. To compute *LoA*_(LD+MAF)_, we start by computing *RA*_Exp_ using fGWAS. For AFR ancestry, *RA*_Exp_ decreases by more than 5 percentage points for BMI, height, HDL-cholesterol, extraversion, subjective well-being, and cannabis use. For most other phenotypes in AFR, values either decrease or remain stable, with the exceptions of educational attainment and drinks per week, which increase. For SAS, substantial decreases (¿5 percentage points) are observed for height, non-HDL cholesterol, migraine, and subjective well-being. In contrast, increases of similar magnitude occur for age at first birth, nearsightedness, and drinks per week. For EAS, most phenotypes show higher *RA*_Exp_ values compared to *RA*_Exp_ estimated using standard-GWAS. Increases of more than 5 percentage points are observed for age at first birth, nearsightedness, morning person, and depressive symptoms, while only height and migraine show substantial decreases. Across all ancestries, only three phenotypes—educational attainment, nearsightedness, and drinks per week—consistently exhibit increases in *RA*_Exp_. Among these, drinks per week shows particularly pronounced gains.

We estimate the *LoA*_(LD+MAF)_ only for phenotypes where *RA*_Obs_ in a non-EUR ancestry was significantly lower than 1 (*p <* 0.05), indicating reduced predictive accuracy relative to the EUR reference group. In AFR, we find *LoA*_(LD+MAF)_ values ranging from 142.19% (S.E. = 16.1%) for non-HDL cholesterol to 61.49% (S.E. = 6.3%) for cigarettes per day; in SAS, from 132.11% (S.E. = 21.5%) for cognitive performance to 16.61% (S.E. = 4.5%) for age at first menses; and in EAS, from 46.74% (S.E. = 12.8%) for age at first birth to 16.59% (S.E. = 5.7%) for hayfever. Estimates with 95% confidence intervals for the full list of phenotypes are reported in Supplementary Table 11.

Relative to standard PGIs, fGWAS-based estimates of *LoA*_(LD+MAF)_ show both upward and downward shifts depending on the ancestry–phenotype combination. In AFR ancestry, *LoA*_(LD+MAF)_ goes down for the majority of phenotypes. In SAS, there are changes in both directions, and in EAS, most *LoA*_(LD+MAF)_ values go up. These contrasts indicate that fGWAS-based PGIs can meaningfully alter the relative contribution of LD and MAF to cross-ancestry prediction loss, with the direction and magnitude of these changes varying by phenotype and ancestry (Figure 8).

**Fig. 8:**
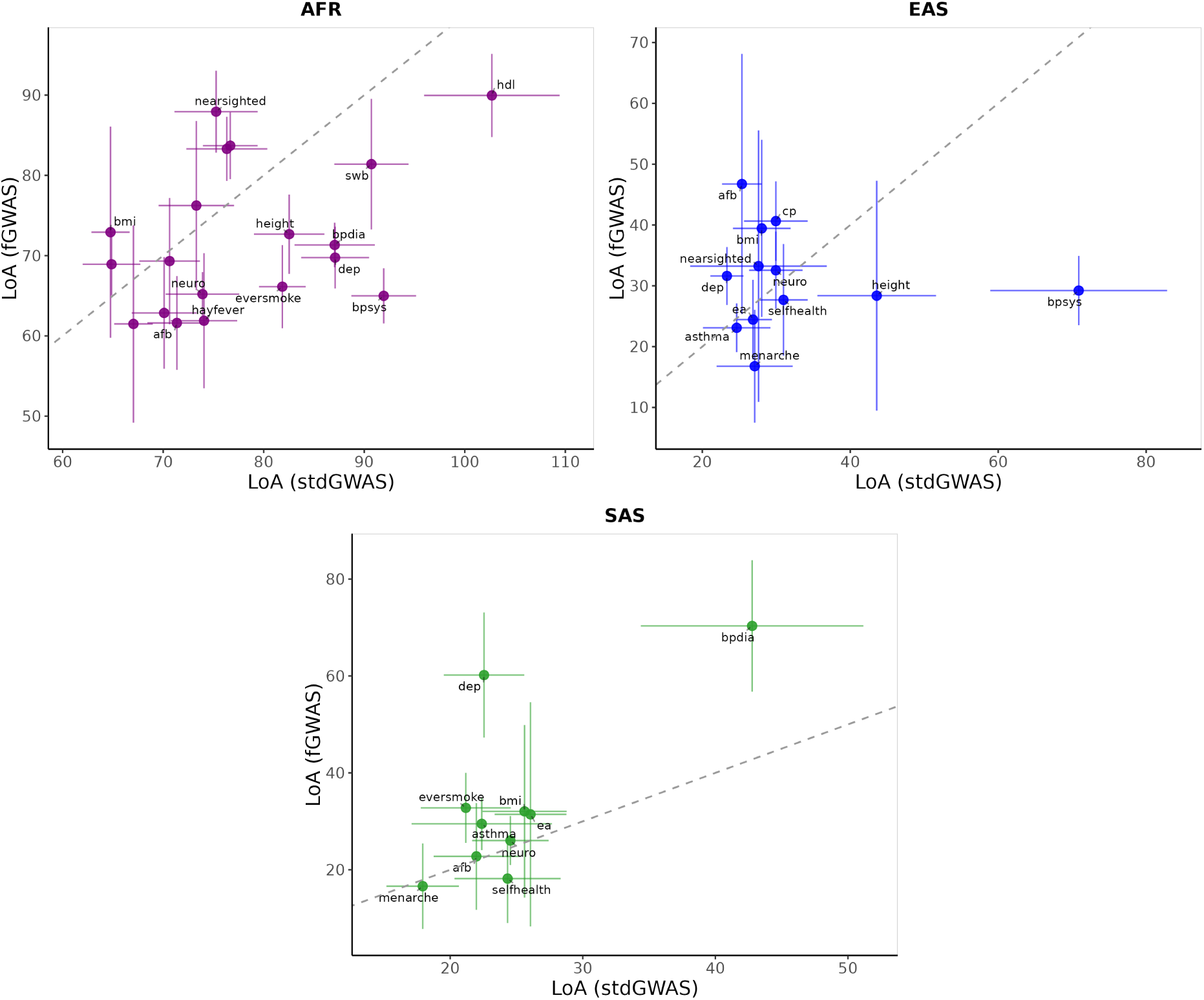
Comparison of *LoA*_(LD+MAF)_ estimates between standard GWAS and family-based GWAS (fGWAS) approaches across non-European genetic ancestries in the UKB cohort. This figure presents three panels, each corresponding to a non-European ancestry (AFR: African, EAS: East Asian, and SAS: South Asian), showing *LoA*_(LD+MAF)_ estimates for individual phenotypes. In each panel, the x-axis represents *LoA*_(LD+MAF)_ derived from standard GWAS-based PGIs, and the y-axis represents estimates from fGWAS-based PGIs, bars are 95% confidence intervals. A dashed 45-degree line in each panel denotes equality between the two estimates. Phenotypes with non-significant shrinkage in *RA*_Obs_ are excluded, as well as those with case proportions *<* 1% (including asthma, breast cancer, prostate cancer, bipolar disorder, and schizophrenia). Additionally, non-HDL cholesterol was omitted for AFR, cognitive performance and drinks per week for SAS, and diastolic blood pressure for EAS, due to imprecise *LoA*_(LD+MAF)_ estimates, but these remain included in Supplementary Tables 5 and 10.

A particularly salient observation is that the *LoA*_(LD+MAF)_ estimate for non-HDL cholesterol in AFR consistently exceeds 100 % with both standard-GWAS and fGWAS-based PGIs—including when heritability is added to the standard-GWAS model—suggesting that the observed predictive accuracy in this subgroup is higher than what LD, MAF, and *h*^2^ differences alone would predict (Figure 8). This persistent observation of *LoA*_(LD+MAF)_ exceeding 100% may reflect inaccuracies in one or more parameters used to compute *RA*_Expected_. For instance, if LD patterns are more similar than assumed, the LD similarity term 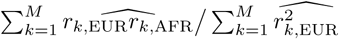 would be underestimated thereby lowering *RA*_Expected_. Overestimation of MAF differences would similarly depress *RA*_Expected_ by inflating the denominator of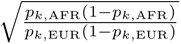. Inaccurate assumptions about effect-size variances—expressed as 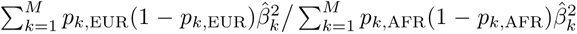—could further bias *RA*_Expected_ downward if causal effects explain more variance in AFR than predicted. These factors collectively may lead to *LoA*_(LD+MAF)_ exceeding 100 %.

## 3 Discussion

In this study, we provide a comprehensive analysis of the factors driving the loss of predictive accuracy of PGIs when PGIs trained on European genetic ancestry samples are applied to other genetic ancestries. Analyzing an extensive set of 54 phenotypes, we expand beyond prior work by comparing the portability of PGIs for biologically proximal traits with more distal behavioral and social traits. Building on the prior framework by Wang *et al.* ^8^, we introduce several methodological advances - including the use of genomewide SNPs, accounting for ancestry-specific heritability estimates, and a comparison of standard versus family-based GWAS PGIs - to provide a more detailed account of PGI portability. Our findings confirm substantial reductions in the predictive accuracy of PGIs for non-European ancestries, with the lowest observed in African, followed by East Asian and South Asian genetic ancestries, consistent with prior studies ^2,8,9,12^. Furthermore, we show that this loss is not uniform across trait categories. Specifically, we find that PGI portability is substantially lower for behavioral and social traits compared to more biologically proximal phenotypes. This pattern likely reflects their greater sensitivity to environmental, cultural, and socio-economic influences, which may interact with genetic effects in population-specific ways.

Our core findings are broadly consistent with Wang *et al.* ^8^, showing that differences in LD and MAF account for a substantial portion of accuracy loss, particularly in African genetic ancestry (83%) compared to much lower contributions in East Asian (34%) and South Asian (25%) ancestries. However, we also observe notable differences in the estimated contribution of these factors for certain traits, which likely stem from methodological distinctions. Our use of genome-wide PGIs, rather than PGIs based only on GWS SNPs, enables a more comprehensive evaluation of polygenic contributions and provides a better benchmark for PGI applications, the majority of which are based on genome-wide PGIs. It also allows us to analyze PGIs for traits that would have too little predictive power if constructed solely from GWS SNPs, enhancing our ability to capture ancestry-specific patterns of predictive accuracy loss for behavioral traits.

Our results for African genetic ancestry align with recent work by Hu *et al.* ^49^, who show widespread conservation of causal effect sizes and conclude that factors like LD and MAF are likely the primary drivers of PGI performance differences. However, our study provides the critical additional insight that this conclusion may not generalize across all non-European populations. The substantially smaller role of LD and MAF in East and South Asian ancestries suggests that other factors—such as cross-ancestry differences in SNP-based heritability or imperfect genetic correlations—are likely the dominant contributors to PGI accuracy loss in these groups.

Another key contribution of our study is the use of fGWAS-based PGIs to assess how PGI portability is affected by the standard-GWAS SNP weights being confounded by passive gene-environment correlation and assortative mating. We find that fGWAS-based PGIs can improve portability for few traits, most notably for BMI in individuals of African genetic ancestry, suggesting that some of the portability gap may be attributable to population-specific biases present in standard PGIs. This finding is in contrast with recent findings by Zhang and Conley ^50^, who reported no improvement in cross-ancestry prediction accuracy in HRS and Add Health when using fGWAS-based PGIs for a number of traits including BMI. However, for the majority of traits analyzed, our findings align with Zhang and Conley ^50^ showing that the relative accuracy between the standard and fGWAS-based approaches is broadly consistent.

Collectively, our findings present an interesting puzzle. While the overall loss of PGI accuracy follows a simple gradient corresponding to genetic distance from the reference group (EUR sample), our analysis suggests that portability of the PGI will vary considerably within a certain genetic ancestry depending on the phenotype. Behavioral phenotypes appear to be less portable across ancestries, and for these phenotypes, the relative contribution of cross-ancestry LD and MAF differences to the loss in predictive accuracy is smaller. Still, cross-ancestry LD and MAF differences appear to explain the majority of accuracy loss in AFR genetic ancestry, where the loss of accuracy is the greatest.

While our study provides valuable insights into the key factors influencing PGI portability, several aspects warrant further investigation. First, our analyses were limited to common SNPs (MAF ¿ 0.01 in each genetic ancestry), which do not fully capture the contribution of rare variants. Rare variants are often population-specific and poorly imputed in reference panels that lack adequate representation of non-European genetic ancestries. Incorporating rare variants and improving imputation strategies for diverse populations could alter the cross-ancestry portability of PGIs. Second, the use of genotype reference panels like the 1000 Genomes which are biased toward European ancestry, may introduce inaccuracies in estimating population parameters such as MAF and LD structure, particularly for non-European ancestries. These inaccuracies could lead to over- or under-estimation of the contribution of various factors, such as LD and MAF, to the loss of PGI predictive accuracy. Increasing the diversity and size of reference panels is crucial for accurate characterization of the genetic architecture of complex traits across genetic ancestries, enabling more robust analyses. Finally, while fGWAS-based PGIs showed promise for improving portability for certain traits, their smaller effective discovery sample sizes and stringent QC filters limit power and genome-wide SNP coverage. Expanding family-based GWAS datasets in scale and coverage could enhance the robustness of fGWAS-based approaches, further advancing our understanding of genetic architecture.

## 4 Methods

### 4.1 Study Cohorts

Our analyses were conducted using two prospective longitudinal studies with genomic data: the UK Biobank (UKB) and the Health and Retirement Study (HRS). Table 1 provides the number of genotyped samples by study cohort and genetic ancestry.

*Discovery Samples:* For this study, we utilized GWAS summary statistics derived from two sources for the standard and family-based GWAS (fGWAS) approaches.

The GWAS summary statistics and PGI weights for the standard GWAS approach were obtained from the PGI-Repository version 2 ^26^, maintained by the Social Science Genetic Association Consortium (SSGAC). For UKB, we used the GWAS summary statistics that were generated for the third partition (UKB3) as defined by the Repository, which consists of one third of the sample after excluding individuals of non-European genetic ancestries and includes only individuals with no third-degree or closer relatives. The summary statistics are based on meta-analyses of GWAS from up to three sources, all of which include only individuals of European genetic ancestries: 23andMe, UKB, and published GWA studies. To avoid sample overlap, discovery meta-analyses for the UKB target cohort excludes the UKB3 partition (leveraging UKB1-UKB2, 23andMe, and published GWAS as applicable), and discovery meta-analyses for the HRS target cohort excludes HRS; Supplementary Table 8 details, by phenotype and cohort, the contributing GWAS sources and the total sample sizes.

The summary statistics for the fGWAS approach were obtained from the largest meta-analyzed fGWAS dataset recently released by Tan *et al.* ^39^. The discovery sample includes up to 16 cohorts and is restricted to individuals of European genetic ancestries. Because neither of our prediction cohorts, UKB and HRS, are family samples, the fGWAS discovery samples do not include them.

*Prediction Samples:* For the standard GWAS-based approach, we used both UKB and HRS as prediction cohorts. For the fGWAS-based approach, only UKB was used as the prediction sample.

In the UKB, the target sample comprised individuals of European genetic ancestries from the third partition (UKB3) and individuals of non-European ancestries. This resulted in a composition of 162,963 individuals of European (EUR) ancestries, 11,413 of South Asian (SAS) ancestries, 2,216 of East Asian (EAS) ancestries, and 9,494 of African (AFR) ancestries. In the Health and Retirement Study (HRS), the target sample included 12,774 individuals of EUR and 3,593 of AFR genetic ancestries. South Asian (SAS, N=87) and East Asian (EAS, N=162) ancestries were excluded from the HRS target sample due to their small sample sizes. Descriptive statistics for all phenotypes in these prediction (target) cohorts are reported in Supplementary Table 12..

### 4.2 Genotyping and Imputation

The details of genotyping and imputation for UKB and HRS can be found in references ^51^ and ^37^, respectively.

### 4.3 Identification of Genetic Ancestries

We follow a PCA-based approach to identify genetic ancestries. To estimate the PCs, in each cohort, we first restricted the genotypes to HapMap3 SNPs ^40^ and converted the dosages to hard calls. Then, we merged these genotypes with the full 1000 Genomes Phase 3 reference sample ^38^, keeping only SNPs that had a call rate *>* 99% and minor allele frequency *>* 1% after the merge. We estimated the loadings for the first 10 PCs in the 1000 Genomes subsample and then projected the remaining samples onto this PC space. We assigned an individual to a genetic ancestry as defined by the 1000 Genomes Project if each of their 10 PCs fell within four standard deviations of the average for that ancestry in the 1000 Genomes sample. Following Wang *et al.* ^8^, we excluded individuals of American genetic ancestry (AMR) due to their complex patterns of genetic admixture.

### 4.4 Construction of PC Controls

We generated ancestry-specific principal components to control for population stratification when estimating the explanatory power (incremental R2) of PGIs. These PCs were obtained using a procedure different than the PCs used for ancestry identification. Prior to generating the PCs, in each ancestry, we removed SNPs meeting any of the following criteria: (1) call rate *<* 99; (2) MAF *<* 0.01; (3) HWE p-value *<* 10*^−^*^5^; (4) imputation accuracy *<* 0.7; (5) SNPs in long-range LD blocks in EUR ancestry (chr5:44mb–51.5 mb, chr6:25mb–33.5 mb, chr8:8mb–12mb and chr11:45mb–57mb). We pruned the remaining SNPs using a 1 Mb rolling window incremented in steps of 5 variants using a cutoff *R*^2^ *<* 0.1. Using these approximately independent variants, we constructed a genomic relatedness matrix in PLINK 1.9 ^52^ to identify pairs of individuals with a relatedness coefficient above 0.05. We excluded one individual from each such pair, estimated the PC loadings for the first 20 PCs in the sample of unrelated individuals that was obtained, and then projected the remaining individuals onto this PC space.

### 4.5 Phenotypes

We started with a set of 61 phenotypes available in the second release of PGI Repository, aggregated into seven categories: biomarkers, anthropometric traits, cognition and education, personality and well-being, health-related traits, fertility and sexual development, psychiatric conditions, and substance use. We analyzed a PGI if the phenotype was available in the target cohort and for binary phenotypes, the case proportion within each genetic ancestry was greater than 1%. 47 phenotypes satisfied these criteria in UKB and 33 in HRS. A full list of phenotypes and relevant inclusion criteria details are provided in Supplementary Tables S1 and S2.

Prior to analysis, we residualized all phenotypes on a set of covariates. If multiple measurements across time were available, we first obtained the standardized residuals from a regression in each wave of the phenotype on sex (unless the phenotype was sex-specific), a second-degree polynomial in age at the time of measurement, and their interactions, and then take the average of these residuals. This averaged phenotype was residualized a second time on the third-degree polynomial in birth year, sex and their interactions. If only a single measurement was available or the phenotype was defined using the maximum recorded value, the phenotype was residualized on sex, a third-degree polynomial in birth year, and their interactions. A more detailed description of phenotype definitions, pre-processing, and handling of repeated measures is provided in Supplementary Table 7.

### 4.6 Estimation of SNP-based heritability

We estimated SNP-based heritability (*h*_SNP_^2^) for each phenotype within each ancestry group in UKB and HRS cohorts using the REML algorithm implemented in BOLT-LMM software (v2.3.4) ^53^. To reduce computational burden, we randomly sampled 50,000 unrelated individuals from the European ancestry group in our UKB estimation sample for this analysis. For all other genetic ancestries in UKB and HRS, we used the full set of unrelated individuals identified through kinship filtering (*π*^ *<* 0.05) using PLINK v1.9 ^54,55^. We restricted the set of SNPs to those present in the HapMap 3 reference panel ^40^ and filtered for MAF ≥ 1%, genotype missingness ≤ 15%, and individual-level missingness ≤ 15%. Phenotypes were residualized prior to analysis on covariates as described in the previous section, except that we additionally inlcuded the first 20 principal components of the genetic relatedness matrix (GRM) and, for UKB only, genotyping batch effects.

### 4.7 Computation of Polygenic Indexes (PGIs)

For the standard GWAS-based approach, we used weights from the second release of PGI Repository to construct the PGIs ^26^, following the same methodology. These weights were obtained for ∼ 2.9 million pruned common variants from the full UKB European-genetic-ancestry (*N* ≈ 450, 000) data set from Lloyd-Jones *et al.* ^25^ by adjusting the estimated effects for LD using the SBayesR methodology implemented in GCTB software ^25,56^. PGIs were computed using PLINK2 ^57^ using genotype dosages. Details on the input GWAS included in the discovery meta-analyses and their respective sample sizes for each phenotype are provided in Supplementary Table 8.

For the fGWAS-based approach, SNP weights were obtained from the Tan *et al.* family-based GWAS ^39^. These weights were adjusted for LD using PRS-CS ^41^ and variants were restricted to HapMap3 SNPs ^40^. The remaining steps were the same as the construction of standard-GWAS based PGIs.

### 4.8 SNP Selection for LD Correlation Analysis

For each phenotype, we started by LD-clumping the GWAS that was used to obtain the weights for the PGI after restricting the set of SNPs to those included in the PGI. The algorithm, implemented in PLINK2 ^57^ starts by selecting the SNP with the lowest association p-value. SNPs within a 2000kb window that are correlated (*R*^2^ *>* 0.01) with the index SNP and had p-values below 0.5 are clumped together with the index SNP. The process iteratively continues by selecting the SNP with the lowest p-value among those that are not yet assigned to a clump and repeating the clumping steps until no SNPs with p-value *<* 0.5 remain. From this list of approximately independent SNPs, we extracted three subsets with the lowest p-values: the top 100, 1,000, and 10,000 SNPs.

Next, we identified SNPs common across the four genetic ancestries (African [AFR], South Asian [SAS], East Asian [EAS], and European [EUR]) from the 1000 Genomes Project Phase 3 reference panel ^38^ after applying the following filters within each genetic ancestry: call rate *>* 95%, MAF *>* 1%, Hardy-Weinberg equilibrium (HWE) p-value *>* 10*^−^*^10^, and subject-level missingness *<* 1%. 4,576,403 SNPs were available in all four genetic ancestries after the filters. We restricted the 1000 Genomes data for each ancestry to this set of SNPs. Then we computed a genetic relatedness matrix using GCTA ^58^ for each ancestry and excluded one individual from each pair that had a relatedness coefficient greater than 0.05.

Then, for each top SNP set (top 100, 1,000, and 10,000 SNPs), we defined candidate causal SNPs as SNPs available in the QC’d 1000 Genomes data ^38^, which are within 100kb of a SNP in the top SNP set and that have (*R*^2^ *>* 0.45) with it.

To compute the LD-related summary statistics required for inferring the LD-correlation parameters in the *RA*_Expected_ formula (Equation 2), we used ldcorpair, a C++ program developed by Wang *et al.* ^8^ and available at their GitHub repository^1^. This program calculates two key LD-based parameters. The first one, 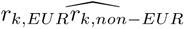, is the average product of the LD between the *k*-th PGI-SNP (i.e. k’th SNP available in the PGI weights) and candidate causal SNPs within a 100kb window of it. The second parameter, 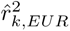, is the mean squared correlation of allele counts between the *k*-th PGI-SNP and the candidate causal SNPs within a 100kb window. We estimated these two parameters for each ancestry using the QC’d 1000 Genomes Project data described above. We repeated the process for each set of 100, 1,000, and 10,000 top SNPs that we identified.

## Data Availability

The GWAS summary statistics and PGI weights used for the standard GWAS-based analyses were obtained from the second release of Social Science Genetic Association Consortium’s (SSGAC) PGI-Repository ^26^ and can be accessed through the SSGAC data portal at https://thessgac.com/ unless the sample includes 23andMe. SNP-level summary statistics from analyses based entirely or in part on 23andMe data can only be reported for up to 10,000 SNPs. Therefore, if the GWAS for a phenotype includes 23andMe, we report summary statistics for only the genome-wide significant SNPs from that analysis. In addition, we report summary statistics for all SNPs from a version of the GWAS that excludes 23andMe. Researchers at non-profit institutions can obtain access to the genome-wide summary statistics from 23andMe used in this paper by completing the 23andMe Publication Dataset Access Request Form, available at https://research.23andme.com/dataset-access/. Family-based GWAS (fGWAS) analyses were conducted using summary statistics from Tan *et al.* ^39^, available through the SSGAC data portal at https://thessgac.com/. For the fGWAS-based analyses, summary statistics for certain phenotypes—blood pressure (diastolic), educational attainment, neuroticism, height, BMI, HDL cholesterol, blood pressure (systolic), depressive symptoms, and non-HDL cholesterol— are based on meta-analyses excluding the HUNT cohort due to their data sharing restrictions.

## Code Availability

All code used for data processing, analysis, and figure generation in this study will be made publicly available upon publication.

## Acknowledgments

This study was supported by Open Philanthropy and the National Institute on Aging/National Institutes of Health through grants R24-AG065184, R01-AG081518 and R01-AG083379. This research was conducted using the UK Biobank Resource under Application Number 11425 and the Health and Retirement Study under Application Number 10535.

## Author Contributions

P.T. and A.O. designed and oversaw the study. R.A. and A.O. performed phenotype and genotype data preparation. R.A. conducted data analysis with guidance and assistance from A.O. and P.T. R.A., A.O., P.T., D.J.B., and A.S.Y. wrote the manuscript.

## 5 Supplementary Information

### 5.1 Estimation of Standard Errors and Confidence Intervals

#### 5.1.1 Incremental *R*^2^ and Observed Relative PGI Predictive Accuracy

We employed a bootstrap procedure with 1,000 replications to estimate the standard errors and construct 95% confidence intervals for incremental *R*^2^ and *RA*_Obs_. This method helps to approximate the sampling distribution of these estimators, assuming that our sample adequately represents the population. In each replication, a new sample was drawn with replacement from the original dataset, and *R*^2^ and *RA*_Obs_ were recalculated using the the analytical models and approaches described in in subsection 2.1. For the construction of 95% confidence intervals, we determined the lower and upper bounds from the 2.5th and 97.5th percentiles of the bootstrap estimates, respectively. This method provides an interval that is expected to contain the true values of *R*^2^ and *RA*_Obs_ with 95% probability, assuming the bootstrap replicates reflect the true variability of these estimators.

#### 5.1.2 Expected Relative PGI Predictive Accuracy

To estimate the standard errors for the expected relative PGI predictive accuracy when *RA*_Expected_ is defined as a function of just MAF and LD, we implemented a leave-one-chromosome-out jackknife approach. This method involves sequentially excluding each of the 22 autosomal chromosomes from the dataset and using the remaining chromosomes to infer relevant population parameters from the 1000 Genomes Project Reference Panel, specifically the minor allele frequencies (MAF) for the candidate causal SNPs and PGI-SNPs, and the mean correlations of allele counts between PGI-SNPs and all candidate causal SNPs within a 100 kb window of the top SNPs. These population parameters and the SNP effect size estimates of the PGIs (PGI-SNPs) were then utilized to compute the *RA*_Expected_ as outlined in Equation 2. After excluding a chromosome and recalculating *RA*_Expected_ based on the inferred parameters, we assessed the variation in *RA*_Expected_ across all iterations to compute the standard error:

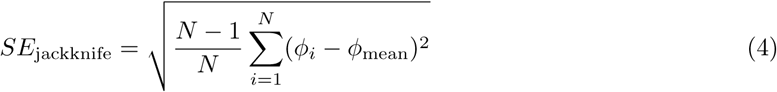

where *N* represents the total number of chromosomes (22), *ϕ_i_* is the estimate from the *i*-th iteration with one chromosome omitted, and *ϕ*_mean_ is the mean of these estimates. This approach provides a robust measure of the standard error by reflecting the stability of *RA*_Expected_ against the exclusion of any single chromosome’s data.

When *RA*_Expected_ is modeled as a function of MAF, LD and *h*^2^, it is calculated by multiplying Equation 2 with the ratio of the SNP-based heritability of the target (non-EUR) to the reference (EUR) genetic ancestries. The standard error of *RA*_Expected_ that accounts for all the three factors is estimated by propagating the errors associated with estimating the respective SNP-based heritabilities using the deltamethod ^48^.

#### 5.1.3 Factors Explaining Loss of PGI Predictive Accuracy

To estimate the loss in predictive accuracy of PGIs attributable to MAF and LD, we calculate *LoA*_(LD+MAF)_ as outlined in Equation (3). We applied the delta method ^48^ to estimate the variability in *LoA*_(LD+MAF)_. This method uses a Taylor series expansion to approximate the variance of functions of random variables. Here, the variance of the loss estimate is influenced by the partial derivatives with respect to *RA*_Expected_ and *RA*_Obs_, calculated as follows:

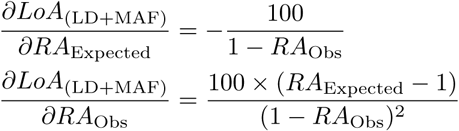

These derivatives and the observed variances of *RA*_pred_ and *RA*_obs_ are then used to estimate the variance of *LoA*_(LD+MAF)_ as follows:

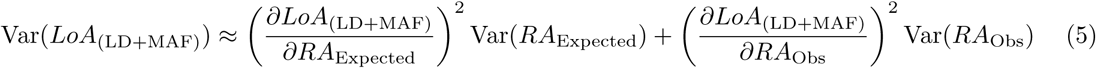

From the variance, the standard error of *LoA*_(LD+MAF)_ is calculated as:

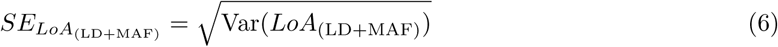

The delta method relies on the assumption that *RA*_Expected_ and *RA*_Obs_ are statistically independent. This assumption is justified in our analysis as these parameters are estimated from non-overlapping samples. This approach allows us to effectively quantify the uncertainty surrounding our estimates of *LoA*_(LD+MAF)_.

### 5.2 Statistical tests

#### 5.2.1 Testing differences in *RA*_Obs_ between fGWAS and standard PGIs

For each ancestry–phenotype, we tested the null hypothesis that the mean difference in observed relative accuracy between fGWAS and standard PGIs equals zero (i.e., *RA*_Obs, fGWAS_ − *RA*_Obs, stdGWAS_ = 0). We used a *paired* nonparametric bootstrap for inference: within each bootstrap replicate, we recomputed *RA*_Obs_ for fGWAS and standard PGIs on the same resampled data and took their difference, thereby constructing an empirical distribution of the difference without imposing a parametric null**^?^**. We chose a nonparametric approach because *RA*_Obs_ is a <u>ratio</u> measure whose sampling distribution can be skewed, heavy-tailed, and heteroskedastic (e.g., when denominators are small or estimates are near zero), making normality-based approximations unreliable. The paired bootstrap preserves the dependence between fGWAS and standard estimates within each resample and naturally accommodates asymmetry.

Two-sided 95% confidence intervals for the difference were obtained via the percentile method (2.5th and 97.5th percentiles of the bootstrap distribution). We also reported empirical, two-sided *p*-values from the same distribution and controlled the false discovery rate across ancestry–phenotype tests using the Benjamini–Hochberg procedure ^42^. A difference was deemed statistically significant when its 95% confidence interval excluded zero and the FDR-adjusted *p*-value met the stated threshold.

#### 5.2.2 Testing differences in *LoA*_(LD+MAF)_ between UKB and HRS in AFR

To test whether the contribution of LD and MAF to the loss of PGI predictive accuracy differs between UKB and HRS in the AFR genetic ancestry, we compared *LoA*_(LD+MAF)_ estimates for each phenotype using a two-sided *Z*-test on the log-transformed values. The log transformation was used to stabilize variance and facilitate interpretation on the ratio scale.

For each cohort and phenotype, we computed the standard error of the log-transformed estimate using the delta method:

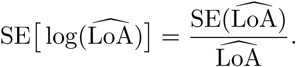

Given that the two cohorts are independent, the standard error of the difference in log-transformed values was:

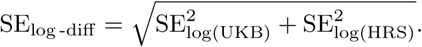

The resulting *Z*-statistic was:

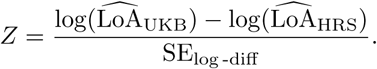

Two-sided *p*-values were obtained from the standard normal distribution, and adjusted for multiple testing using the Benjamini–Hochberg procedure ^42^. A phenotype was considered to show a statistically significant difference in *LoA*_(LD+MAF)_ between cohorts if the FDR-adjusted *p*-value was below 0.05.

---

**Fig. S1:**
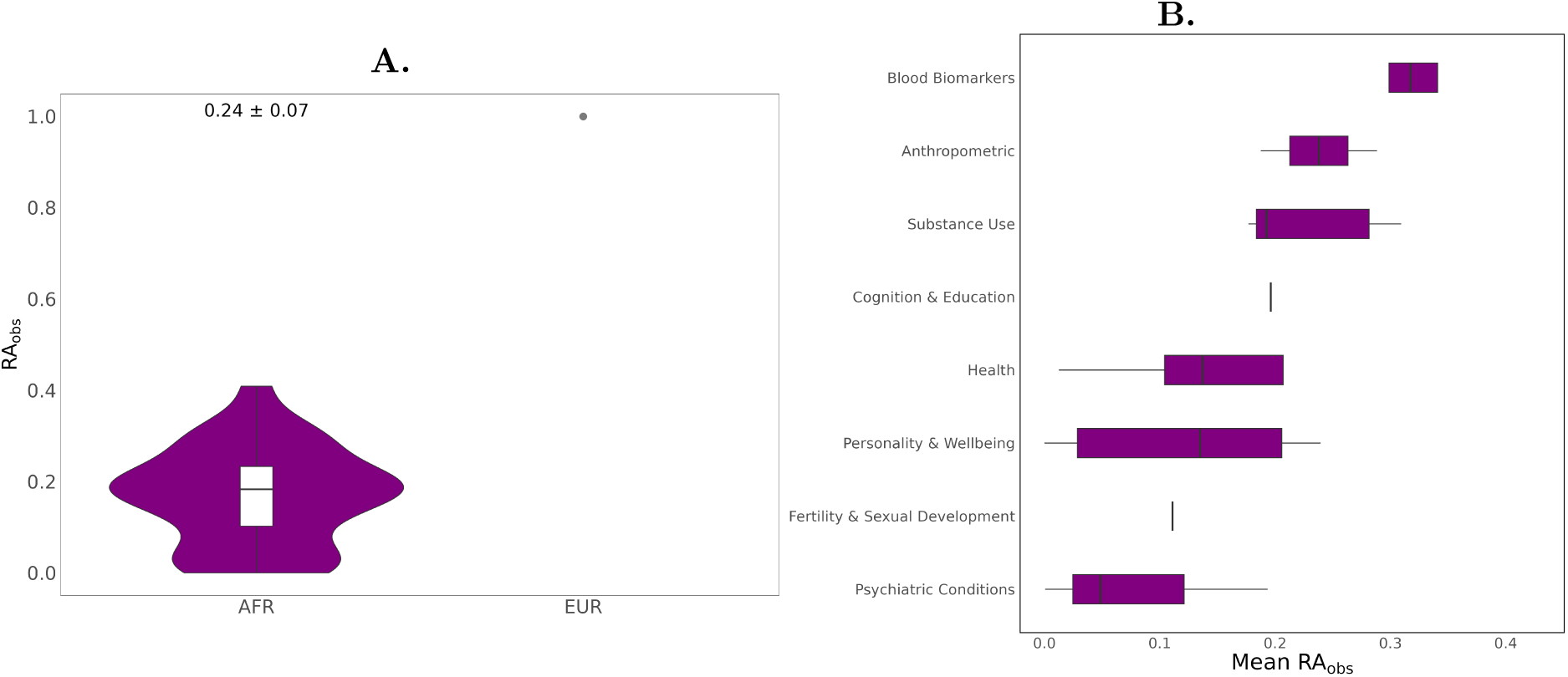
Observed relative predictive accuracy of PGIs in African genetic-ancestry samples from the HRS cohort. **(A)** Violin plot illustrating the distribution of observed relative predictive accuracy (*RA*_Obs_) for PGIs across all phenotypes, with an embedded boxplot representing the quartiles. A grey dotted line at *RA*_Obs_ = 1 represents the EUR genetic-ancestry reference group. The average *RA*_Obs_ and standard error, estimated via a leave-one-phenotype-out jackknife approach, are displayed above the violin plot. **(B)** Breakdown of *RA*_Obs_ by eight phenotype categories. Phenotype exclusions, as described in Figure S2, apply here as well, omitting asthma, breast cancer, prostate cancer, bipolar disorder, and schizophrenia due to case proportions below 1%. Moreover, the phenotype alcohol misuse was also removed from the plots as the estimate is too imprecise, distorting the plot (however, all *RA*_Obs_ are reported in the supplementary tables).

**Fig. S2:**
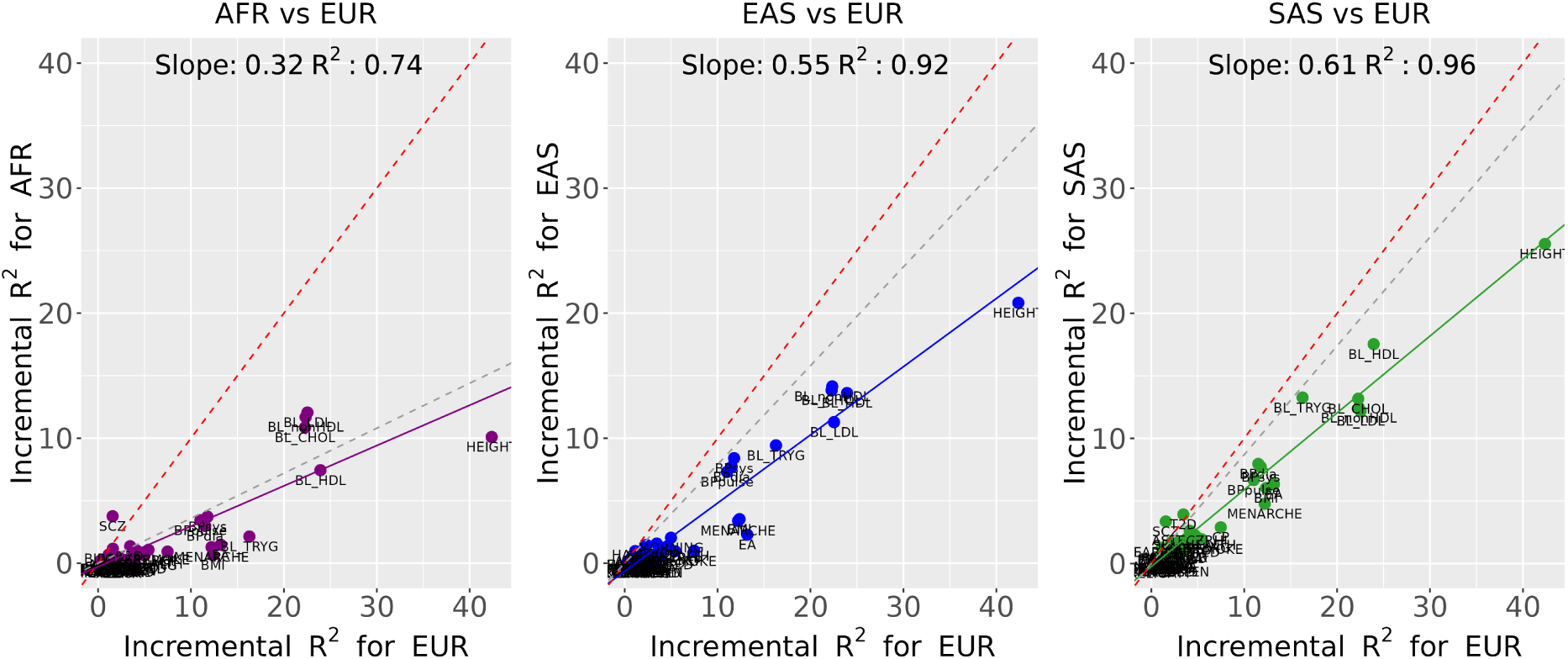
Predictive accuracy of polygenic indexes (PGIs) in non-European versus European genetic-ancestry samples in the UKB cohort. Each panel shows a scatter plot of incremental *R*^2^ values for PGIs in one of three non-European ancestries (AFR: African, EAS: East Asian, SAS: South Asian) compared to EUR (European) incremental *R*^2^ values across various phenotypes. The x-axis in each panel represents the incremental *R*^2^ of a phenotype in EUR, while the y-axis represents the incremental *R*^2^ of the corresponding phenotype for a given non-EUR ancestry. The first panel shows AFR vs. EUR, the second shows EAS vs. EUR, and the third shows SAS vs. EUR. Only phenotypes with an expected *R*^2^ ≥ 1% in EUR ancestry were included, and binary phenotypes with case proportions of less than 1% in the UKB cohort (partition 3) sample—specifically inflammatory bowel disease (IBD), anorexia nervosa, attention deficit and hyperactivity disorder (ADHD), and autism spectrum disorder—were excluded. Points represent individual phenotypes, with colors corresponding to ancestry pairs (AFR: purple, EAS: blue, SAS: green). The solid lines, colored to match each non-EUR ancestry, indicate the fitted linear regression line for each ancestry pair, with the slope and *R*^2^ of this line displayed in the upper right of each panel. The red dashed line represents a slope of 1 (indicating equal predictive accuracy across ancestries), while the secondary gray dashed line denotes the average expected predictive accuracy of PGIs i.e. (*RA*_Expected_) that accounts only for minor allele frequency (MAF) and linkage disequilibrium (LD) differences in each non-EUR genetic ancestry (AFR: 0.36, EAS: 0.79, SAS: 0.88).

**Fig. S3:**
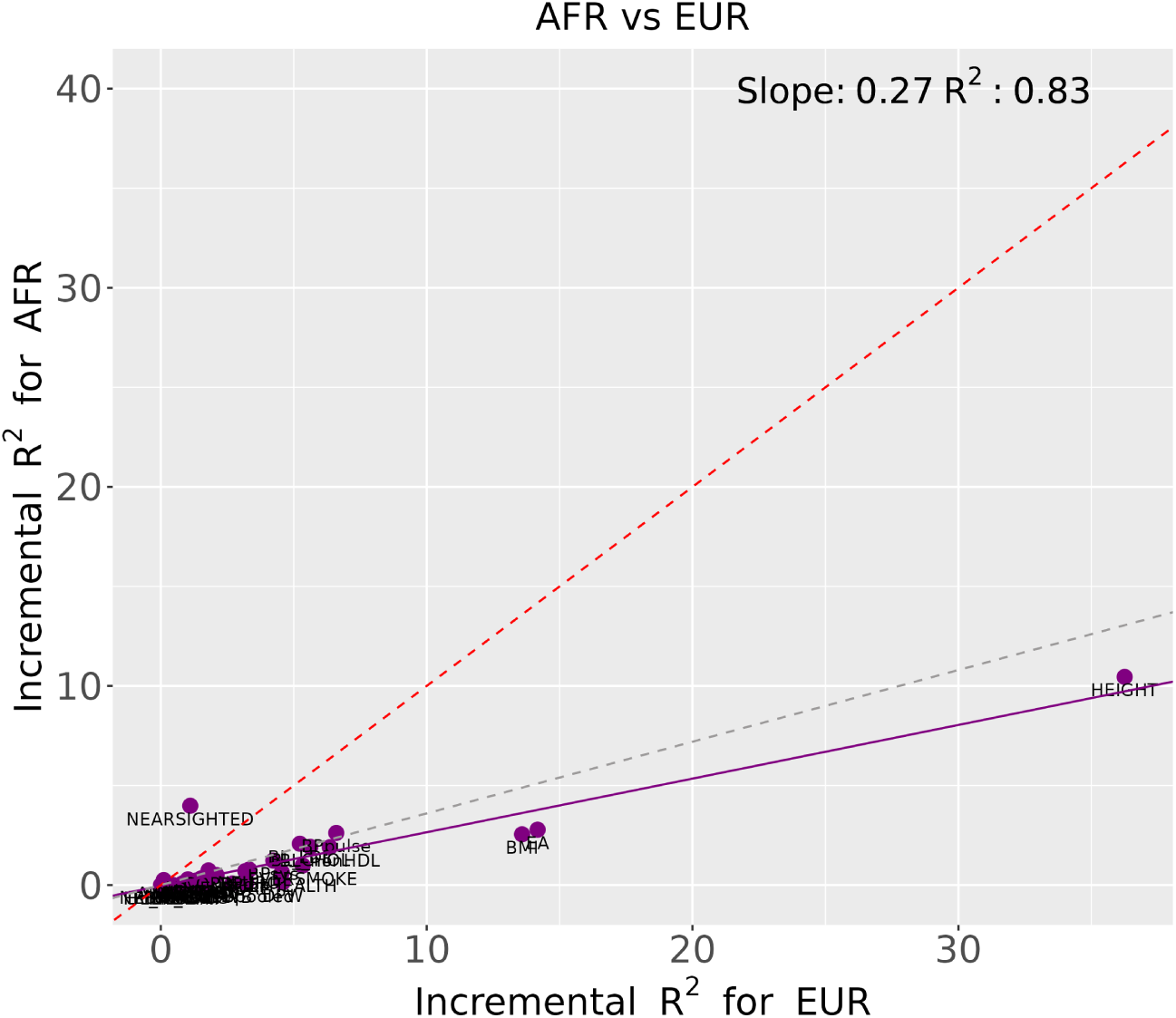
Predictive accuracy of polygenic indexes (PGIs) in African versus European genetic-ancestry samples in the HRS cohort. This panel compares incremental *R*^2^ values for PGIs in AFR (African) versus EUR (European) genetic ancestries across various phenotypes in the HRS cohort. Details follow those described in Figure S1, with binary phenotypes excluded if their case proportions were below 1% in HRS (asthma, breast cancer, prostate cancer, bipolar disorder, and schizophrenia). Points represent individual phenotypes, with a solid regression line fitted for AFR and reference lines indicating the expected predictive accuracy (*RA*_Expected_) based on MAF and LD differences.

**Fig. S4:**
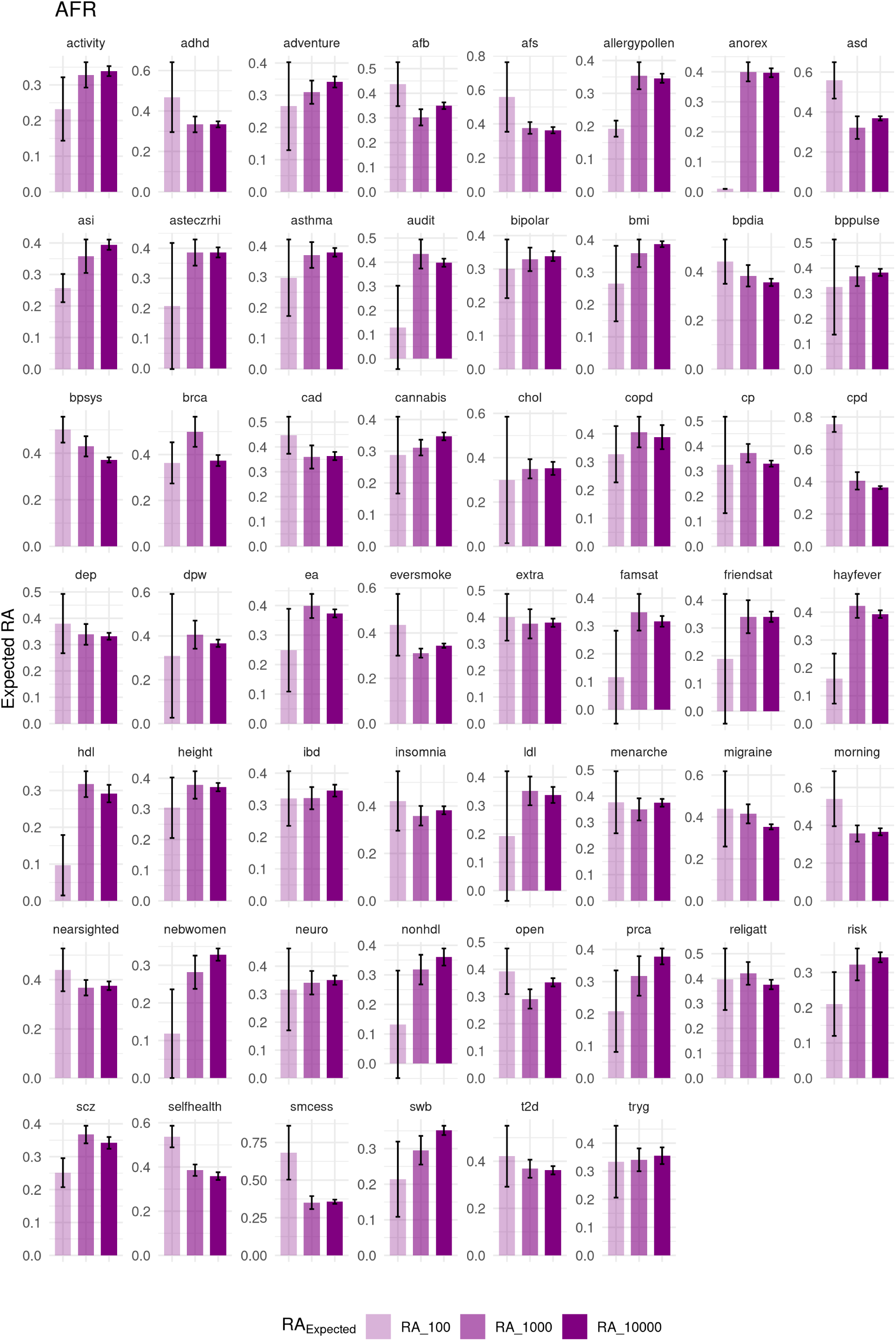
Expected relative accuracy of PGIs in AFR genetic ancestry by top SNP sets in the UKB cohort.

**Fig. S5:**
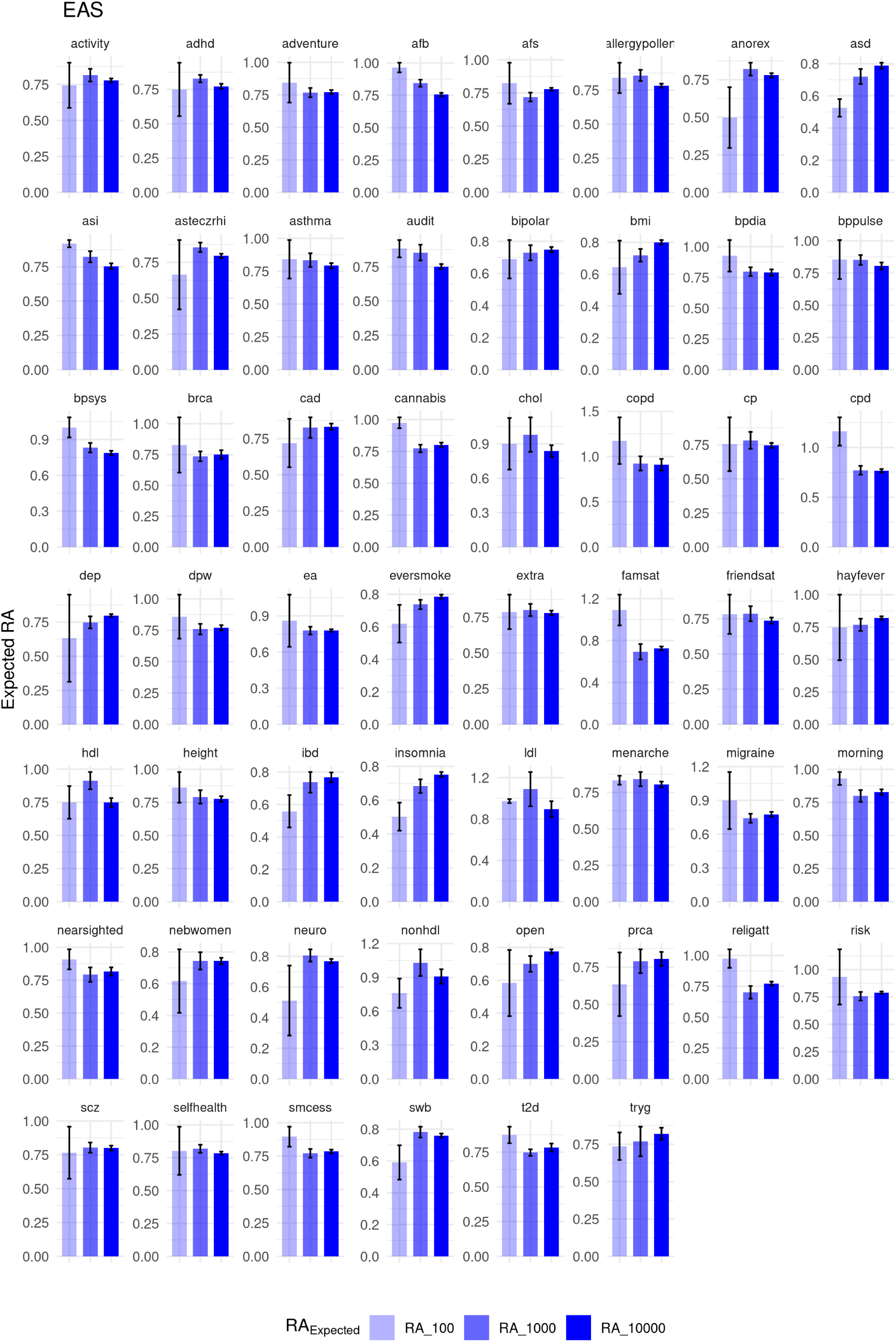
Expected relative accuracy of PGIs in EAS genetic ancestry by top SNP sets in the UKB cohort.

**Fig. S6:**
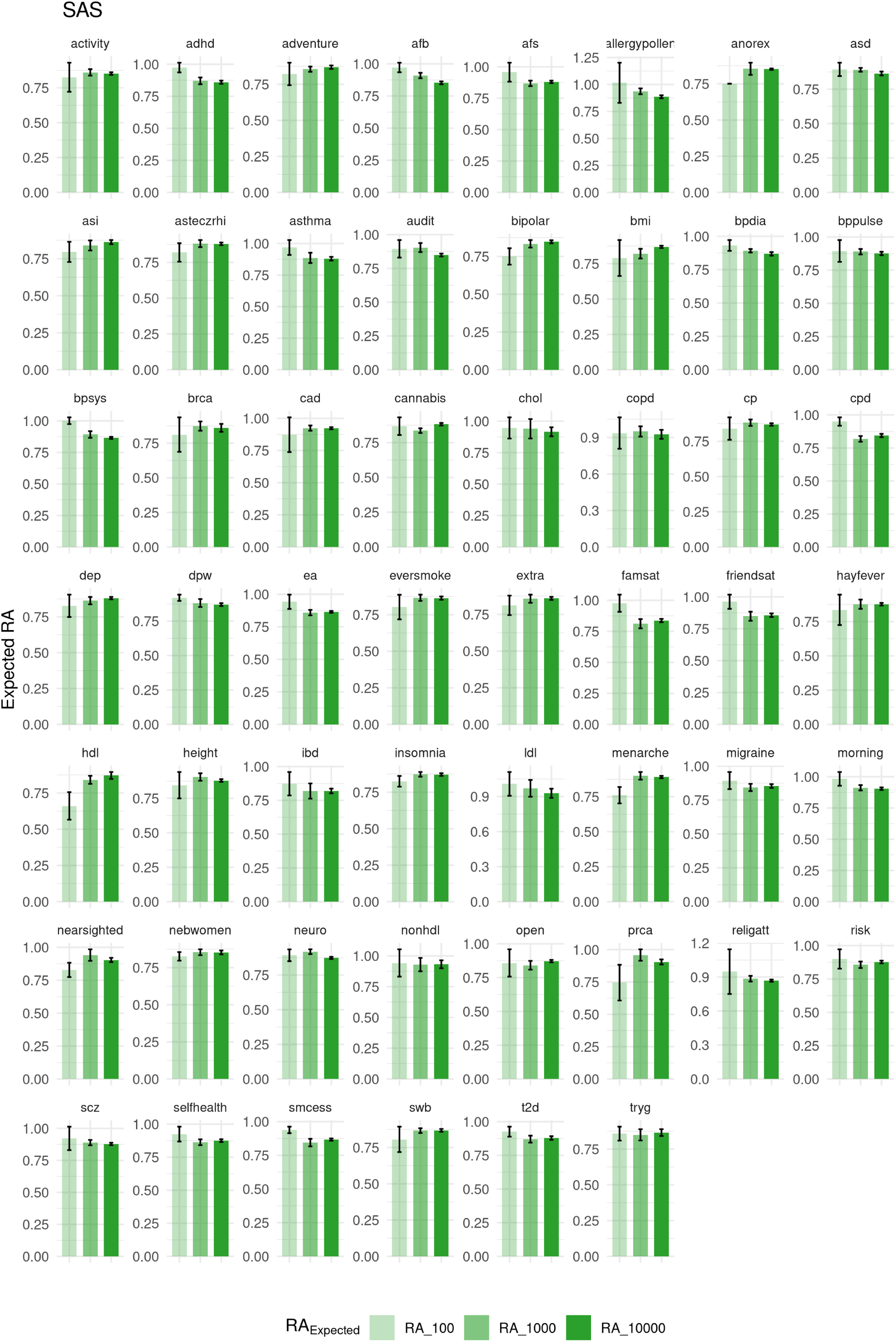
Expected relative accuracy of PGIs in SAS genetic ancestry by top SNP sets in the UKB cohort.

**Fig. S7:**
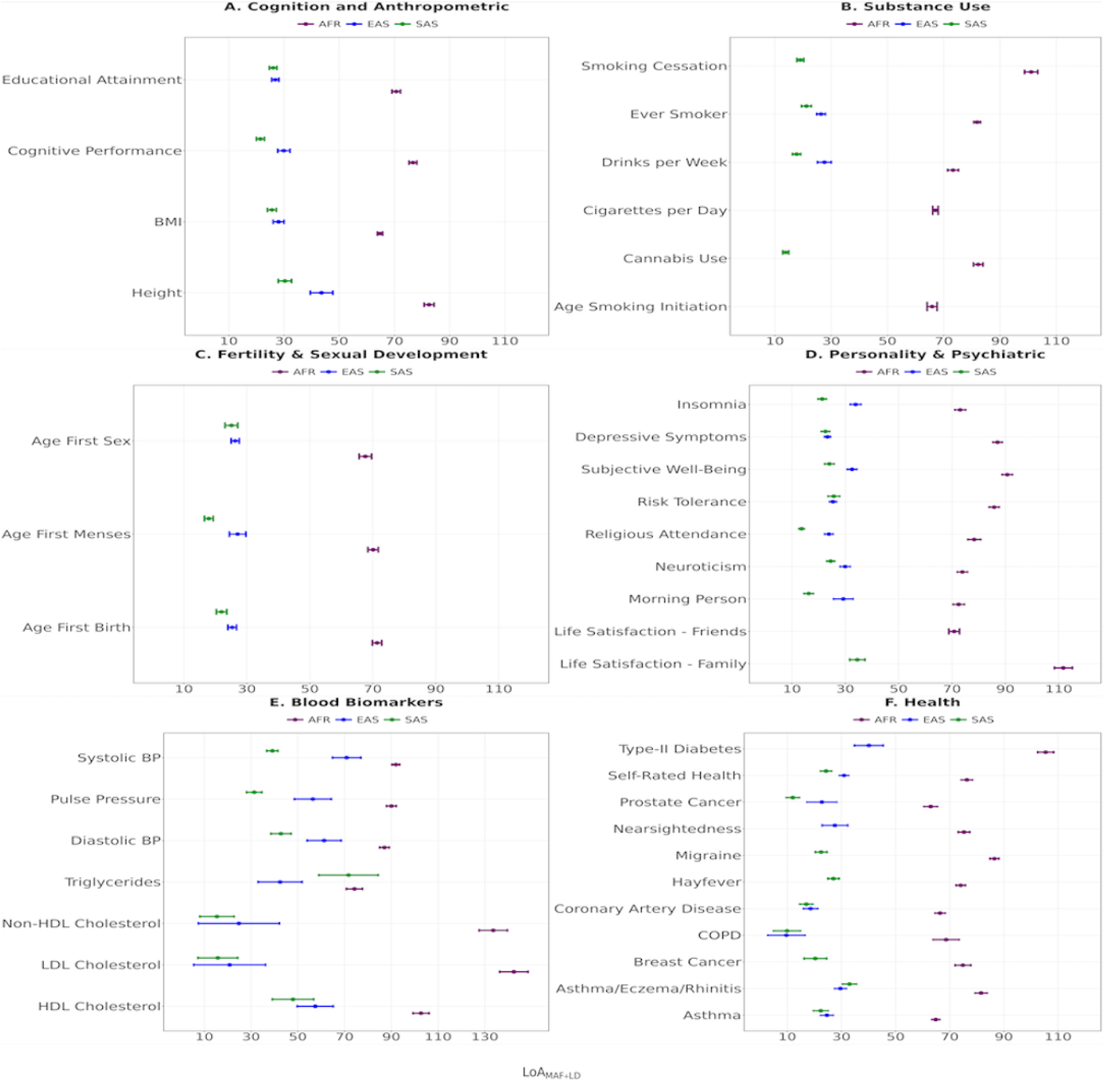
Contribution of Minor Allele Frequency (MAF) and Linkage Disequilibrium (LD) to the Loss of PGI Predictive Accuracy by Genetic Ancestry and Phenotype Category in the UKB Cohort. This figure presents the contribution of MAF and LD to the loss of PGI predictive accuracy (*LoA*_(LD+MAF)_) across <u>six</u> phenotype categories in the UKB cohort: cognition and anthropometric traits (Panel A), substance use (Panel B), fertility and sexual development (Panel C), personality and psychiatric traits (Panel D), blood biomarkers (Panel E), and health (Panel F). For each category, *LoA*_(LD+MAF)_ is shown as point estimates with 95% confidence-interval error bars for the three non-European ancestries—South Asian (SAS), East Asian (EAS), and African (AFR). The *x*-axis indicates *LoA*_(LD+MAF)_, and the *y*-axis lists the phenotypes within each category. Standard errors for *LoA*_(LD+MAF)_ were computed via the delta method, accounting for variance in both *RA*_Expected_ and *RA*_Obs_ in Equation (3) (see Supplementary Note 5.1.3). Binary phenotypes excluded due to low case proportions (*<* 1%) in UKB are inflammatory bowel disease (IBD), anorexia nervosa, attentiondeficit/hyperactivity disorder (ADHD), and autism spectrum disorder (ASD).

**Fig. S8:**
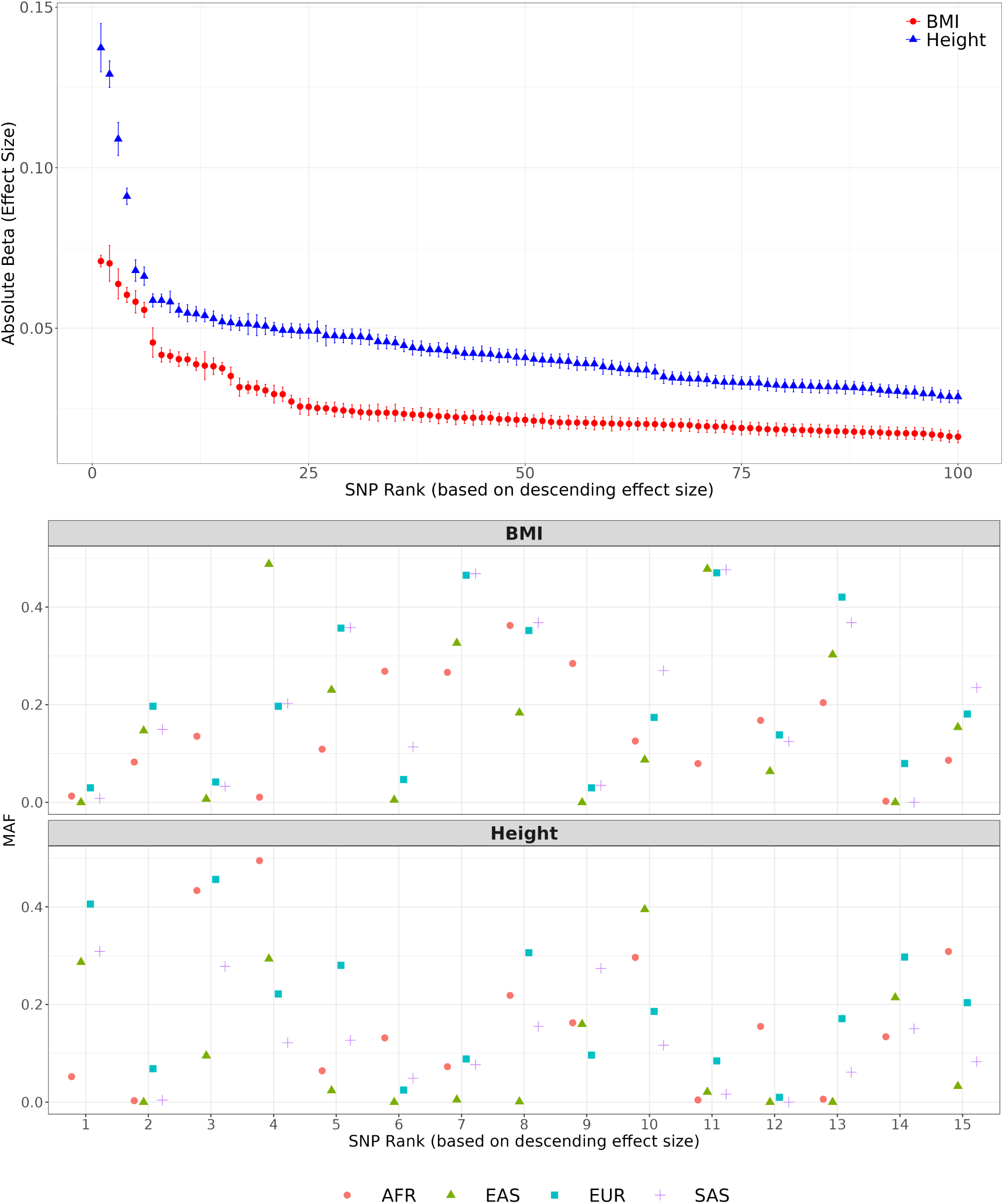
Effect sizes and minor allele frequencies of SNPs associated with anthropometric traits from standard GWAS. The top 100 SNPs were selected based on their p-values and are ordered by descending absolute effect size. **Top panel:** Absolute effect sizes of these top 100 SNPs, ranked accordingly. **Bottom panel:** Minor allele frequencies (MAF) for the top 15 large-effect SNPs disaggregated by genetic ancestry (EUR, SAS, AFR, and EAS) as estimated from the 1000 Genomes Project (1KGP) reference panel ^38^. MAF values were computed as the minimum of the reported alternate allele frequency and its complement (i.e. MAF = min(ALT FREQ, 1 *−* ALT FREQ)).

**Fig. S9:**
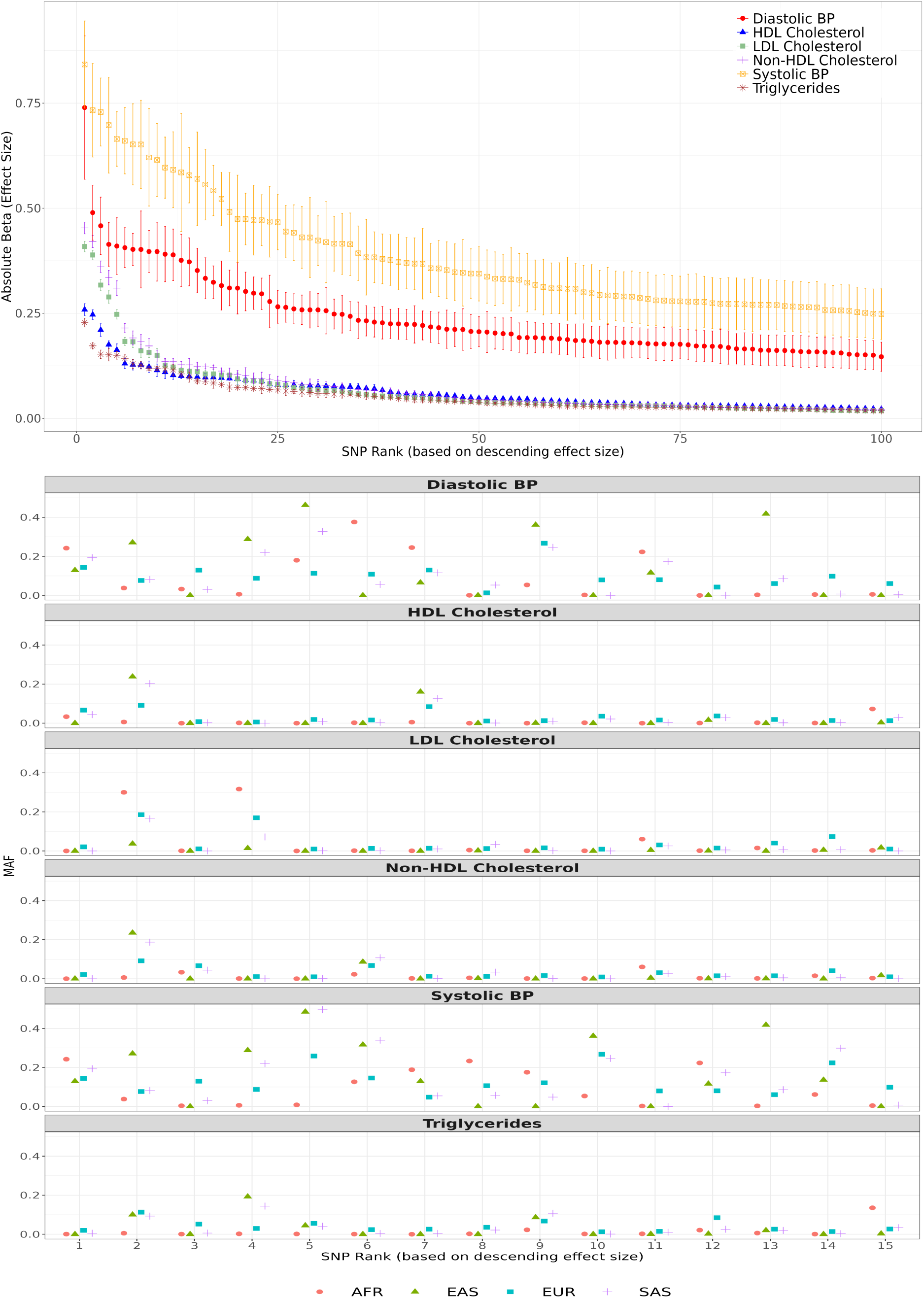
Effect sizes and minor allele frequencies of SNPs associated with blood biomarker traits from standard GWAS. The top 100 SNPs were selected based on their p-values and are ordered by descending absolute effect size. **Top panel:** Absolute effect sizes of these top 100 SNPs, ranked accordingly. **Bottom panel:** Minor allele frequencies (MAF) for the top 15 large-effect SNPs disaggregated by genetic ancestry (EUR, SAS, AFR, and EAS) as estimated from the 1KGP reference panel ^38^. For both panels, MAF is computed as the minimum of the reported alternate allele frequency and its complement (i.e. MAF = min(ALT FREQ, 1 *−* ALT FREQ)).

**Fig. S10:**
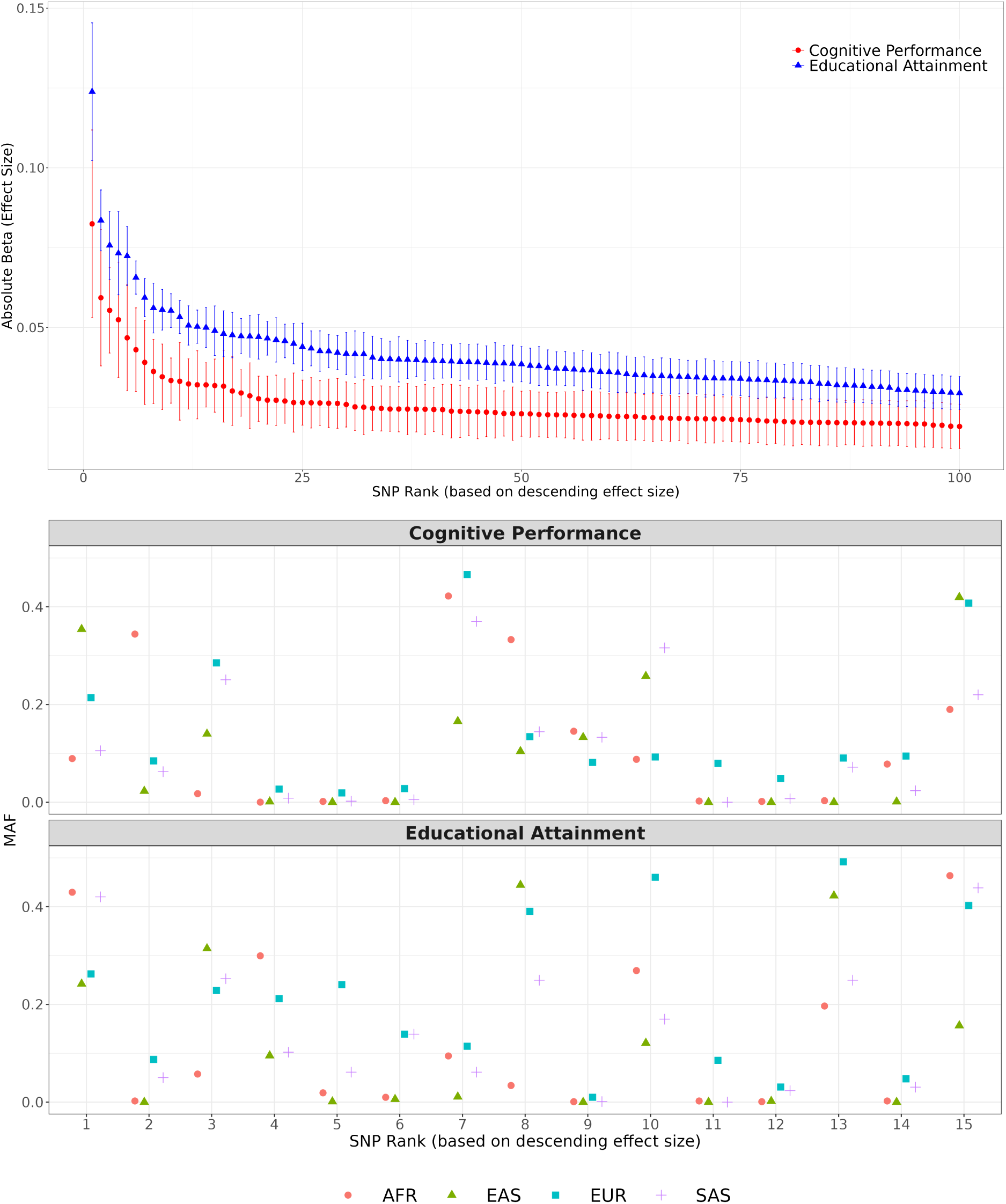
Effect sizes and minor allele frequencies of SNPs associated with cognition and education traits. The top 100 SNPs were selected based on their p-values and are ordered by descending absolute effect size. **Top panel:** Absolute effect sizes of these top 100 SNPs, ranked accordingly. **Bottom panel:** Minor allele frequencies (MAF) for the top 15 large-effect SNPs disaggregated by genetic ancestry (EUR, SAS, AFR, and EAS) as estimated from the 1KGP reference panel ^38^. For both panels, MAF is computed as the minimum of the reported alternate allele frequency and its complement (i.e. MAF = min(ALT FREQ, 1 *−* ALT FREQ)).

**Fig. S11:**
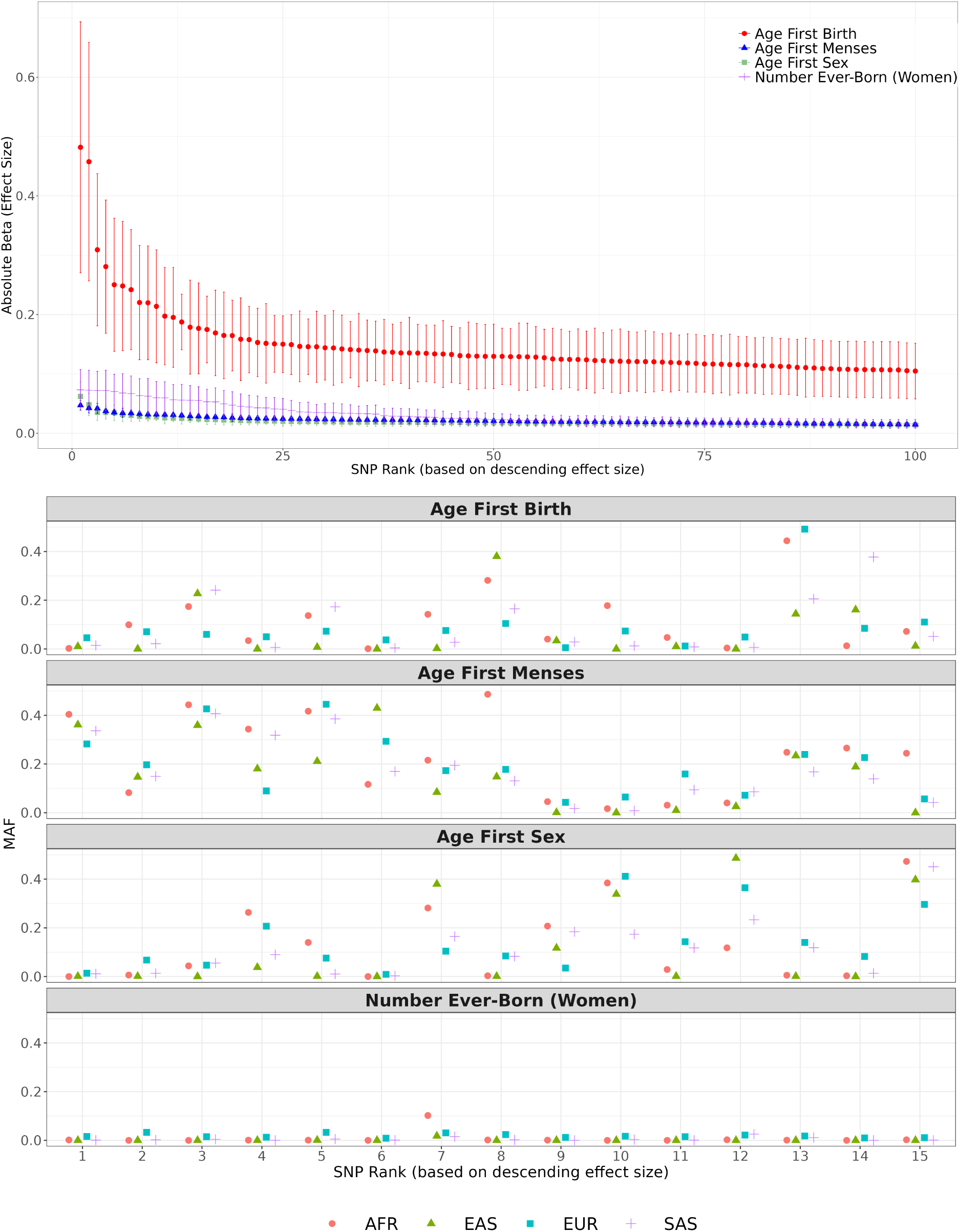
Effect sizes and minor allele frequencies of SNPs associated with fertility and sexual development traits. The top 100 SNPs were selected based on their p-values and are ordered by descending absolute effect size. **Top panel:** Absolute effect sizes of these top 100 SNPs, ranked accordingly. **Bottom panel:** Minor allele frequencies (MAF) for the top 15 large-effect SNPs disaggregated by genetic ancestry (EUR, SAS, AFR, and EAS) as estimated from the 1KGP reference panel ^38^. For both panels, MAF is computed as the minimum of the reported alternate allele frequency and its complement (i.e. MAF = min(ALT FREQ, 1 *−* ALT FREQ)).

**Fig. S12:**
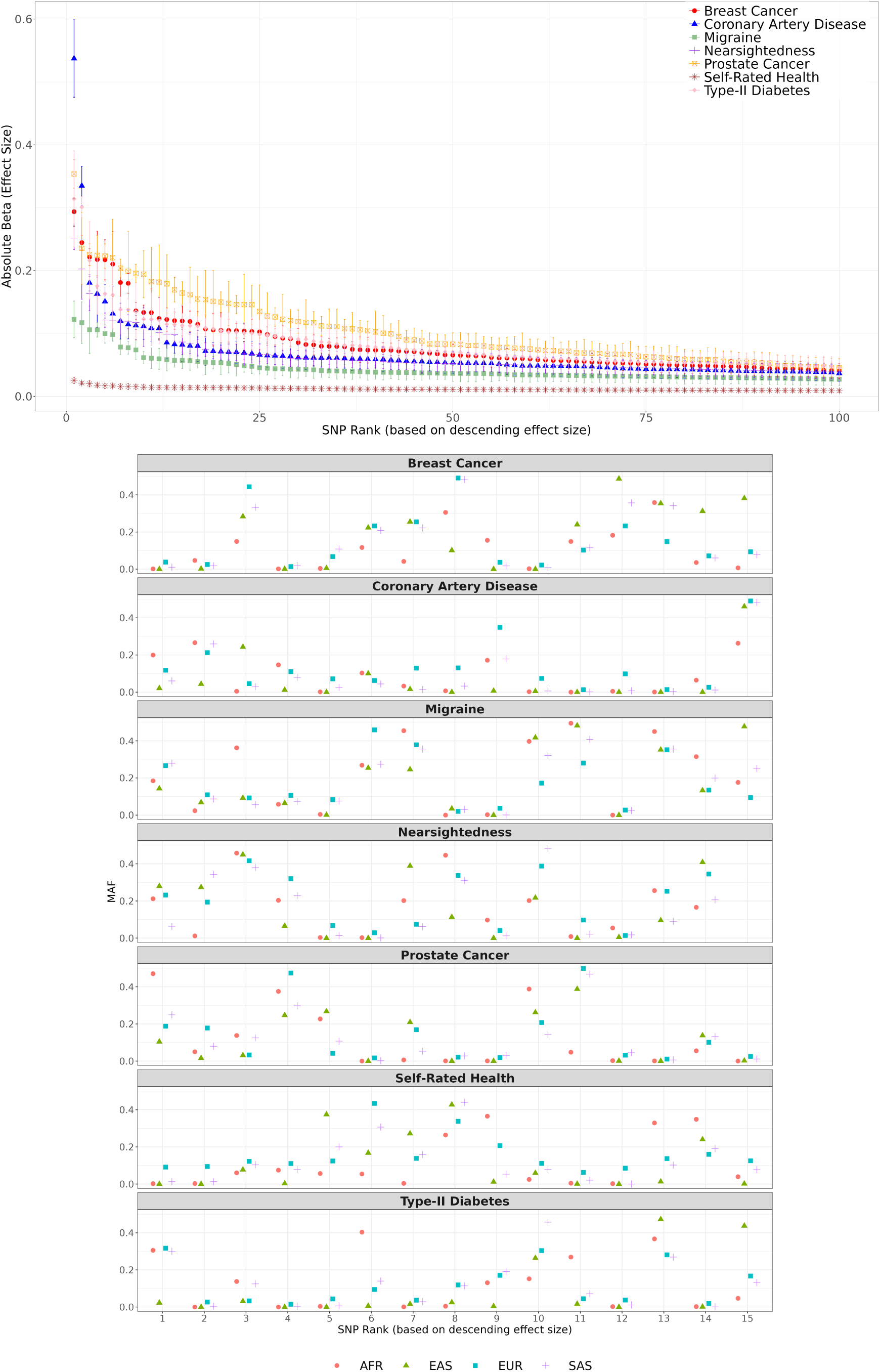
Effect sizes and minor allele frequencies of SNPs associated with health-related traits (other than respiratory and allergic conditions). The top 100 SNPs were selected based on their p-values and are ordered by descending absolute effect size. **Top panel:** Absolute effect sizes of these top 100 SNPs, ranked accordingly. **Bottom panel:** Minor allele frequencies (MAF) for the top 15 large-effect SNPs disaggregated by genetic ancestry (EUR, SAS, AFR, and EAS) as estimated from the 1KGP reference panel ^38^. For both panels, MAF is computed as the minimum of the reported alternate allele frequency and its complement (i.e. MAF = min(ALT FREQ, 1 *−* ALT FREQ)).

**Fig. S13:**
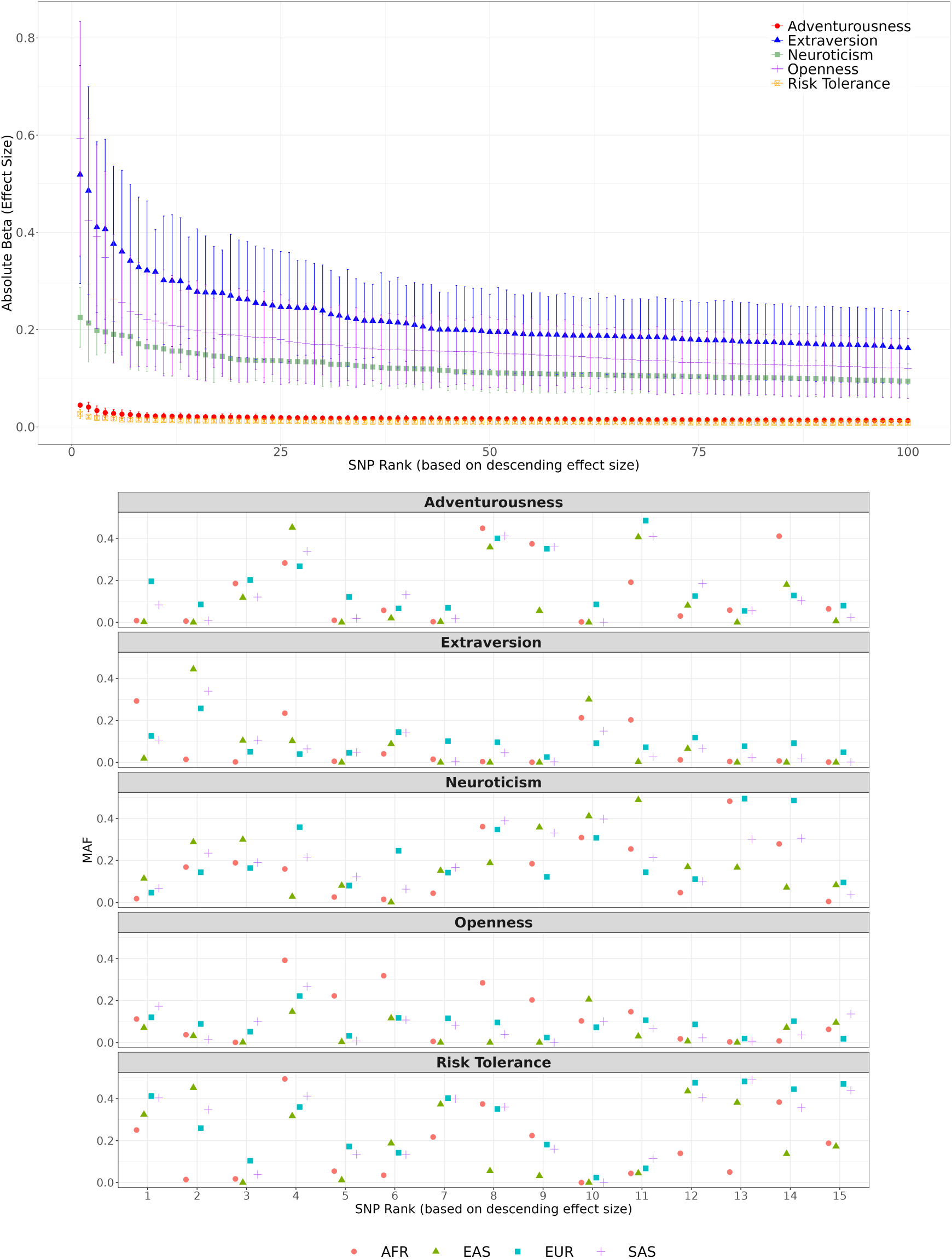
Effect sizes and minor allele frequencies of SNPs associated with personality traits. The top 100 SNPs were selected based on their p-values and are ordered by descending absolute effect size. **Top panel:** Absolute effect sizes of these top 100 SNPs, ranked accordingly. **Bottom panel:** Minor allele frequencies (MAF) for the top 15 large-effect SNPs disaggregated by genetic ancestry (EUR, SAS, AFR, and EAS) as estimated from the 1KGP reference panel ^38^. For both panels, MAF is computed as the minimum of the reported alternate allele frequency and its complement (i.e. MAF = min(ALT FREQ, 1 *−* ALT FREQ)).

**Fig. S14:**
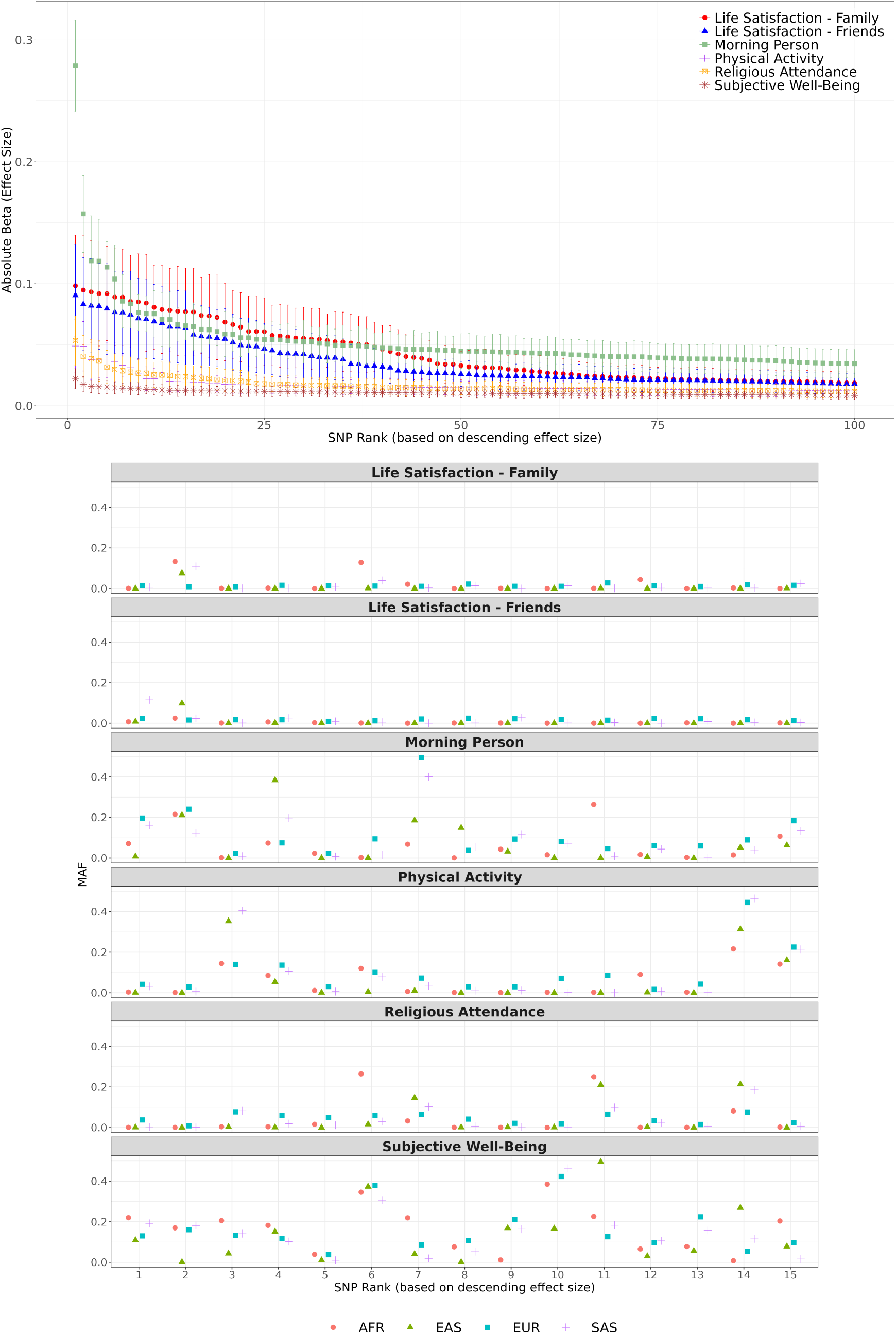
Effect sizes and minor allele frequencies of SNPs associated with wellbeing and lifestyle traits. The top 100 SNPs were selected based on their p-values and are ordered by descending absolute effect size. **Top panel:**

**Fig. S15:**
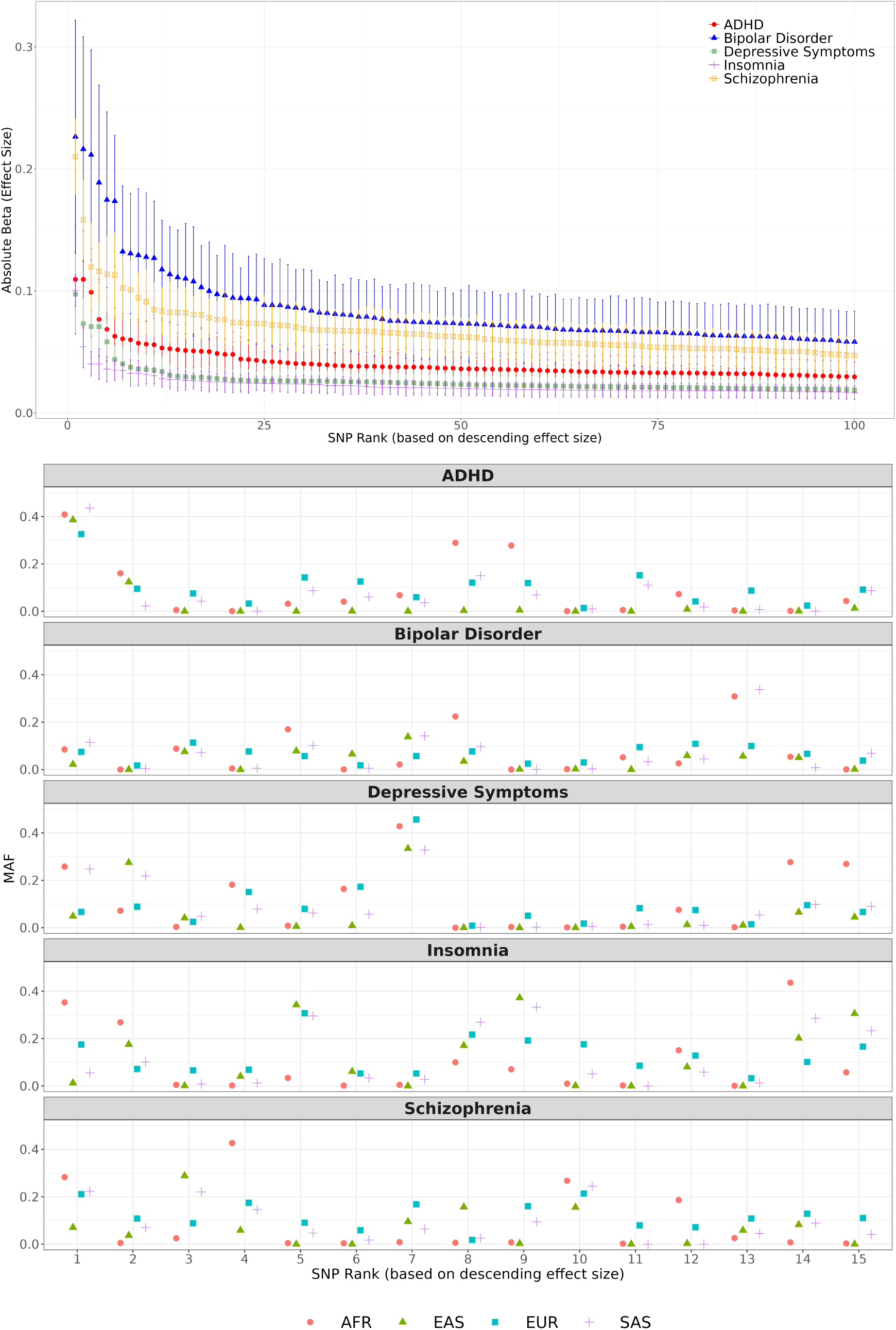
Effect sizes and minor allele frequencies of SNPs associated with psychiatric conditions. The top 100 SNPs were selected based on their p-values and are ordered by descending absolute effect size. **Top panel:**

**Fig. S16:**
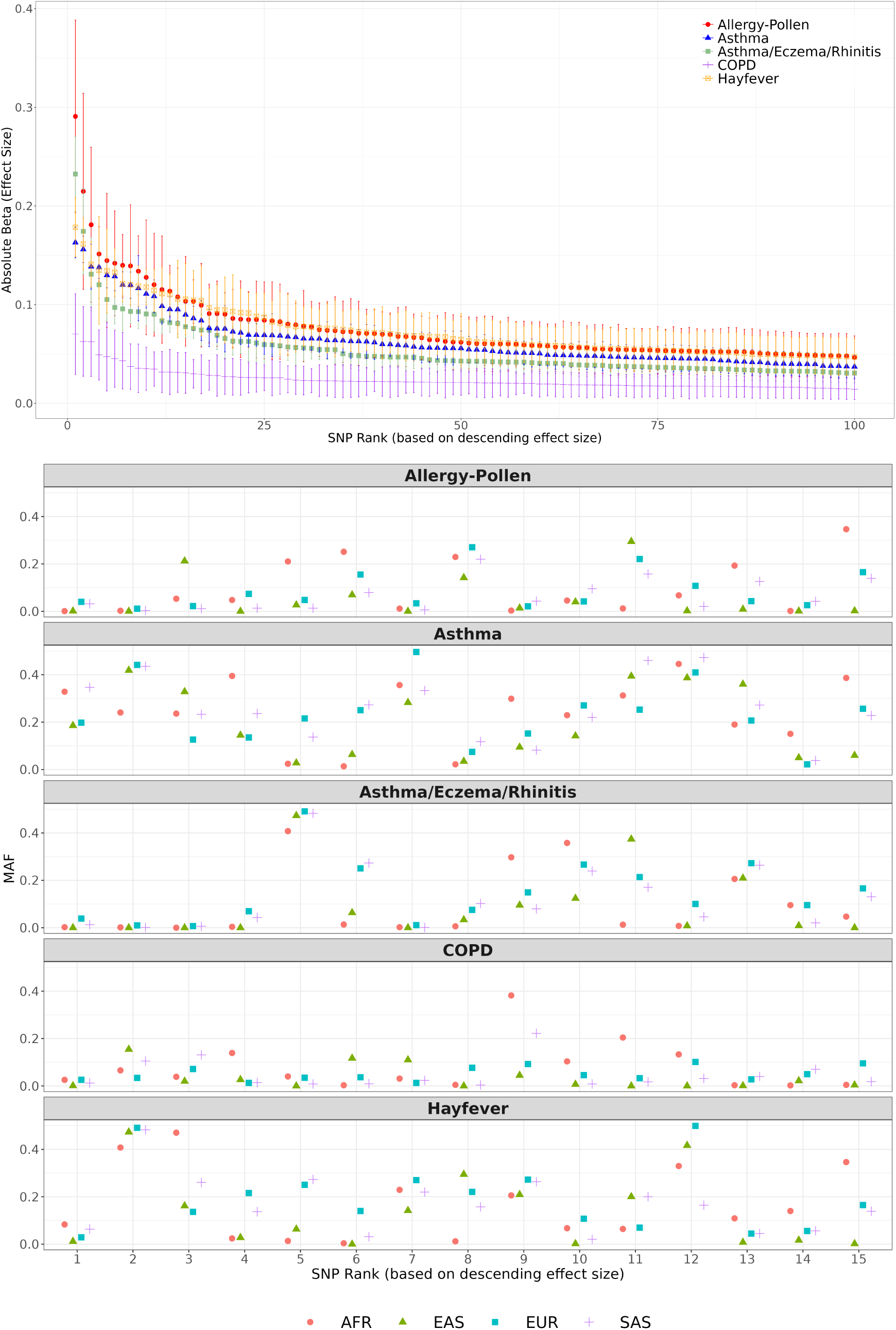
Effect sizes and minor allele frequencies of SNPs associated with respiratory and allergic conditions. The top 100 SNPs were selected based on their p-values and are ordered by descending absolute effect size. **Top panel:** Absolute effect sizes of these top 100 SNPs, ranked accordingly. **Bottom panel:** Minor allele frequencies (MAF) for the top 15 large-effect SNPs disaggregated by genetic ancestry (EUR, SAS, AFR, and EAS) as estimated from the 1KGP reference panel ^38^. For both panels, MAF is computed as the minimum of the reported alternate allele frequency and its complement (i.e. MAF = min(ALT FREQ, 1 *−* ALT FREQ)).

**Fig. S17:**
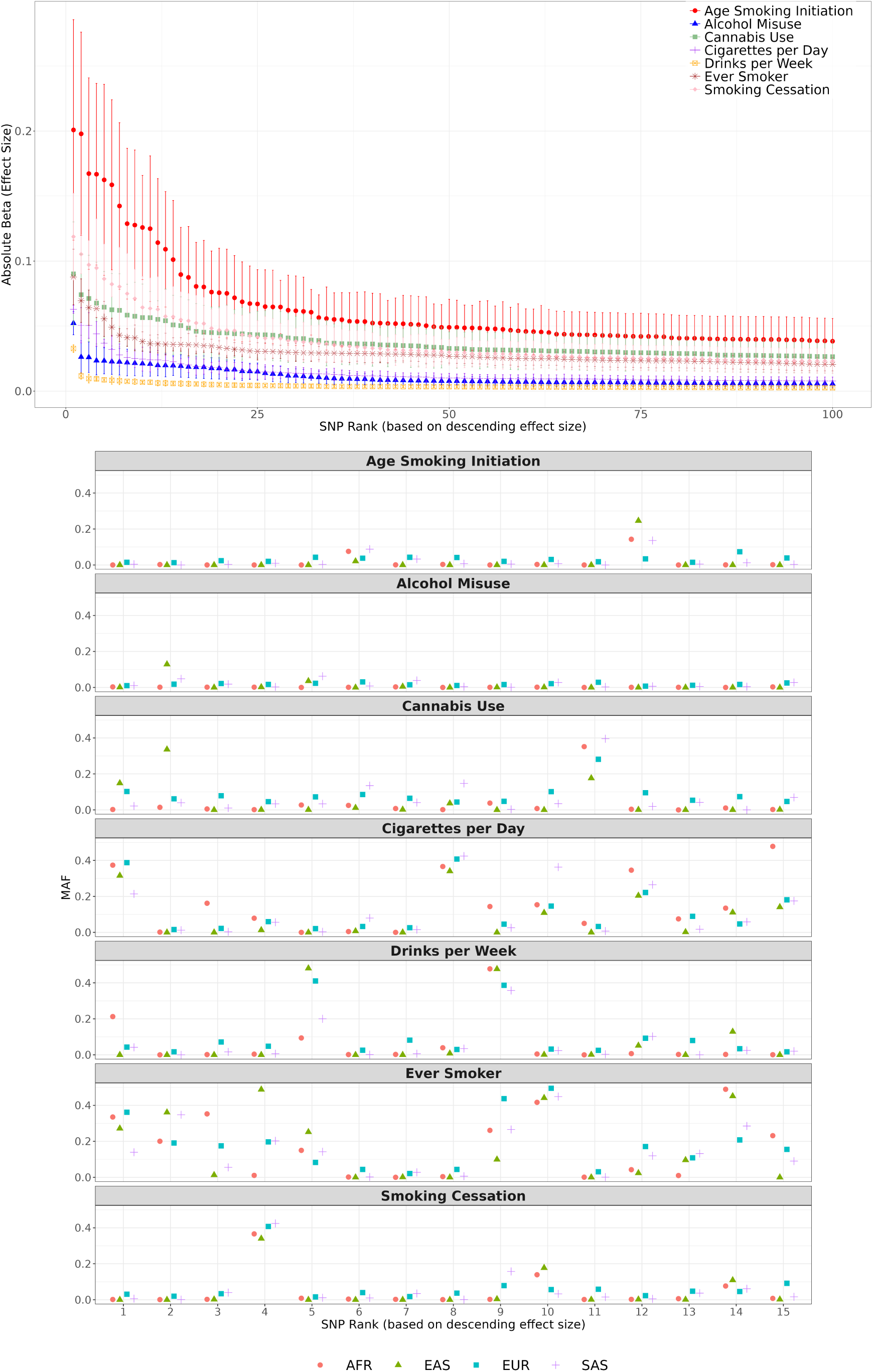
Effect sizes and minor allele frequencies of the top 100 SNPs associated with substance use traits. The top 100 SNPs were selected based on their p-values and are ordered by descending absolute effect size. **Top panel:** Absolute effect sizes of these top 100 SNPs, ranked accordingly. **Bottom panel:** Minor allele frequencies (MAF) for the top 15 large-effect SNPs disaggregated by genetic ancestry (EUR, SAS, AFR, and EAS) as estimated from the 1KGP reference panel ^38^. For both panels, MAF is computed as the minimum of the reported alternate allele frequency and its complement (i.e. MAF = min(ALT FREQ, 1 *−* ALT FREQ)).

## Footnotes

1 https://github.com/loic-yengo/Code for Wang et al2020

